# Bridging immunotypes and enterotypes using a systems immunology approach

**DOI:** 10.1101/2024.11.29.625344

**Authors:** Fabio Affaticati, My K. Ha, Thies Gehrmann, Ilke De Boeck, Maria Kuznetsova, Romi Vandoren, Vincent Van Deuren, Hilde Jansens, Hans De Reu, Jolien Schippers, Karin Peeters, Esther Bartholomeus, Sabrina Van Ierssel, Samuel Coenen, Reinout Naesens, Kevin K. Ariën, Koen Vercauteren, Erika Vlieghe, Philippe Beutels, Pierre Van Damme, Herman Goossens, Eva Lion, Arvid Suls, Sarah Lebeer, Kris Laukens, Pieter Meysman, Benson Ogunjimi

**Affiliations:** Antwerp Unit for Data Analysis and Computation in Immunology and Sequencing (AUDACIS), University of Antwerp; Antwerp, Belgium; ADReM Data Lab, Department of Mathematics and Computer Science, University of Antwerp; Antwerp, Belgium; Center for Health Economics Research and Modelling Infectious Diseases (CHERMID), Vaccine and Infectious Disease Institute, University of Antwerp; Wilrijk, Belgium; Antwerp Center for Translational Immunology and Virology (ACTIV), Vaccine and Infectious Disease Institute, University of Antwerp, Wilrijk, Belgium; Laboratory of Applied Microbiology and Biotechnology, University of Antwerp; Antwerp, Belgium; Department of Clinical Microbiology, Antwerp University Hospital, Antwerp, Belgium; Laboratory of Experimental Hematology (LEH), Vaccine and Infectious Disease Institute, University of Antwerp; Wilrijk, Belgium; Flow Cytometry and Cell Sorting Core Facility (FACSUA), University of Antwerp; Wilrijk, Belgium; Department of General Internal Medicine, Infectious Disease and Tropical Medicine, Antwerp University Hospital; Edegem, Belgium; Laboratory of Medical Microbiology (LMM), Vaccine & Infectious Disease Institute (VAXINFECTIO), University of Antwerp; Wilrijk, Belgium; Center for General Practice, Department of Family Medicine and Population Health (FAMPOP), University of Antwerp; Wilrijk, Belgium; Department of Clinical Biology, Antwerp Hospital Network; Antwerp, Belgium; Virology Unit, Department of Biomedical Sciences, Institute of Tropical Medicine; Antwerp, Belgium; Clinical Virology Unit, Department of Clinical Sciences, Institute of Tropical Medicine; Antwerp, Belgium; Global Health Institute, University of Antwerp; Wilrijk, Belgium; Centre for the Evaluation of Vaccination (CEV), Vaccine and Infectious Disease Institute, University of Antwerp; Wilrijk, Belgium; Centre for Medical Genetics, Edegem, Belgium; Biomedical Informatics Research Network Antwerp (biomina), University of Antwerp; Antwerp, Belgium; Department of Pediatrics, Antwerp University Hospital; Edegem, Belgium

**Keywords:** System immunology, Multi-modal data integration, Clustering analysis, Blood Transcriptomics, Mass cytometry, Inflammation, Immunosenescence, Enterotypes, Adaptive immunity

## Abstract

Unveiling the systemic effects of disease and health requires an holistic approach that has mainly revolved around well established, directly determinable molecular relationships such as the protein synthesis cascade and epigenetic mechanisms. In this study, involving 394 individuals, we found direct linkage of branches spanning human biological functions often not studied in conjunction, using clinical data, gut microbial abundances, blood immune cell repertoires, blood transcriptomic and blood T cell receptor data. Contrary to current paradigms, we demonstrate that immunotypes and enterotypes are orthogonal, likely fulfilling distinct roles in maintaining homeostasis, only bridged via the blood transcriptome. We also identified two distinct inflammatory profiles: the first driven by interferon signalling and the other characterised by non-viral, NF-kB and IL-6 markers. Lastly, we present compelling data showing strong associations of the Ruminococcaceae and Christensenellaceae bacteria with a healthy immunotype and transcriptomic pattern, highlighting their potential role in immune health.

**Graphical Abstract:** 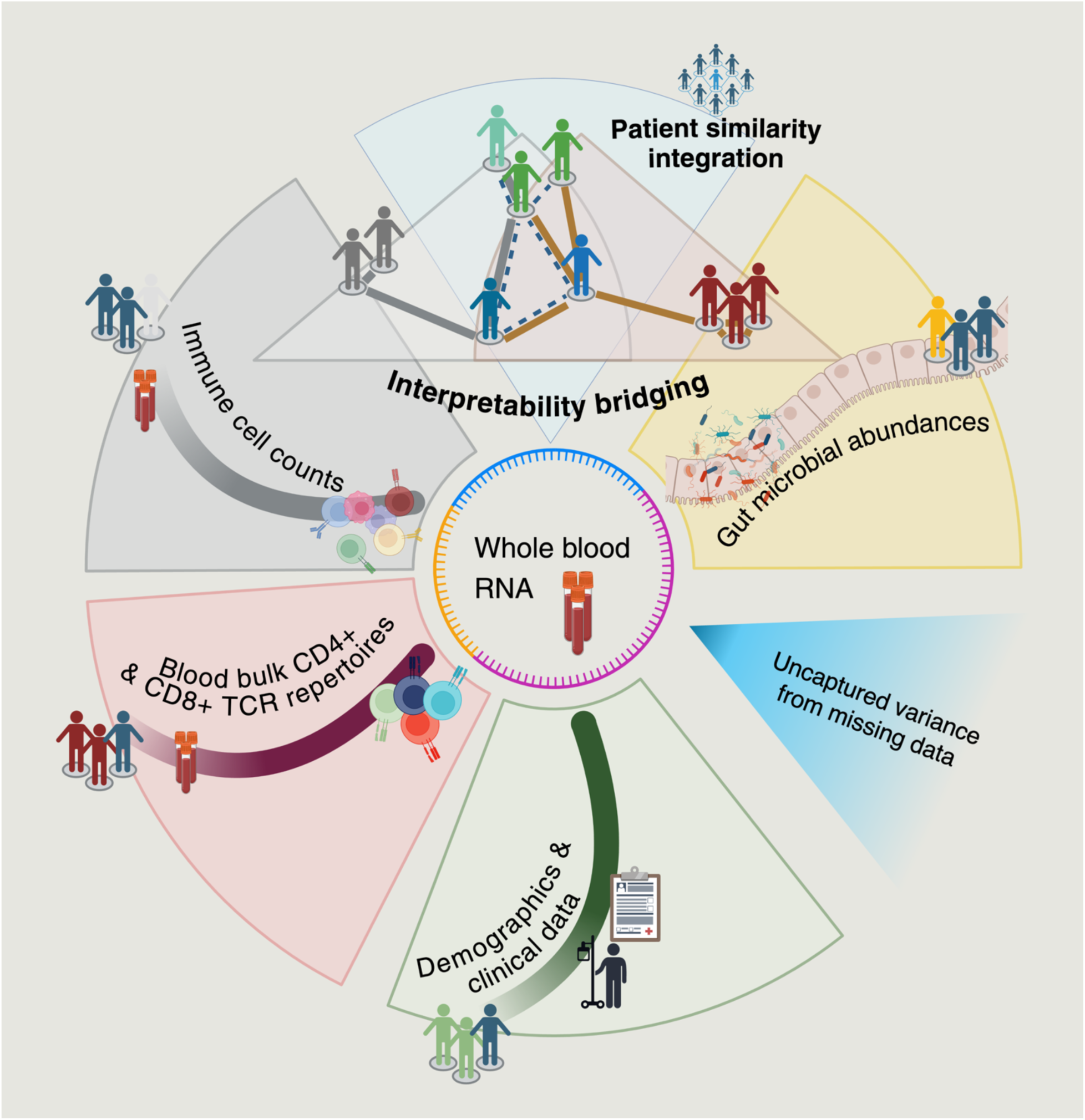

## Introduction

The “immunotype” of an individual can be defined as the broad collection of features that jointly outline their immune condition, ranging from the composition of immune cell populations to diversity and functionality of T-cell receptor (TCR) repertoires. This concept is grounded in the multi-faceted nature of immune status, however, the fundamental question of whether immunotypes manifest as distinct profiles or exhibit continuous variation along a spectrum remains an unsolved puzzle^1^.

The gut microbiome composition is similarly often classified into distinct groups or ‘enterotypes’ ^2^, and it has been theorized that the gut ecosystem has a broad influence on the systemic immune responses, from immune system development^3^ and homeostasis to disease susceptibility^4^. This hypothesis suggests that the gut microbial dynamics may play a pivotal, active role in the establishment of the immunotype, hence participating in a self-regulating feedback loop.

Furthermore, the implications of TCR repertoires for diagnostics and therapeutics are now more apparent than ever, since common patterns within TCR repertoires have been linked to various pathological conditions such as autoimmune diseases, infectious diseases and cancer^5–11^ .

Yet, transitioning from a single data layer to integrating several data modalities hides the complexity of their interactions. Most multi-omics methods have been fully tuned for single-cell technologies, often neglecting valuable bulk data, and around specific datasets such as The Cancer Genome Atlas^12^. Be it simultaneous or sequential integration, these methods often rely solely on the genome-transcriptome-proteome-metabolome axis of well explained interconnections^13^ without modeling further components of the immunotype^14^. Furthermore, while single-cell sampling for large numbers of individuals is cost prohibitive^15^, bulk data, along with clinical information, empowers the generation of sizeable study cohorts and offers a feasible solution to address the challenge.

Within this context, uncovering shared parameters that can capture drivers of fluctuation in the immune status, even in healthy subjects^16^, is imperative and should be at the core of systems immunology research. However, such a setting is wholly understudied due to its complexity and lack of a ground truth to use as anchor point. Shifts from the accepted paradigm of health have recently begun from a passive state to active, adaptive mechanisms of resistance and tolerance^17^.

In this study, we aimed to elucidate if and how different biological modalities are linked, by assembling a multimodal data set from 394 participants presenting at least one data type each. These modalities included peripheral immune cell counts (measured via CyTOF mass cytometry), gut microbial abundances, whole blood RNA sequencing, blood bulk CD4+ and CD8+ TCR repertoires and clinical parameters. These were combined through Similarity Network Fusion for multimodal analysis^18^. Spectral clustering was applied at every step to segregate individuals in homogeneous groups before applying statistical testing to determine characterizing factors of diversity.

We consistently assessed significant differences in T cell exhaustion and senescent markers, observed a *Ruminococcaceae* enterotype association with naivety of the adaptive immune system, characterized clusters based on cytomegalovirus seropositivity, age and BMI. Additionally, deregulated inflammation-related transcriptomic modules served as a guide for interpretation of the study’s findings.

## Results

### Blood CyTOF profiles identify immunosenescence, viral associated and T cell exhaustion markers as basis for multimodal analyses

CyTOF offers a robust way to quantify immune cell phenotypes through mass cytometric staining of cellular markers^19^. Here, we profiled 42 markers simultaneously from 236 individuals (as shown in Figure S1, Table S1 and S2) and subsequently processed the data using FlowSOM^20^. Non-parametric testing of the distributions of these markers allowed us to identify emerging groups. A clustering of the CyTOF data (Materials and Methods) revealed six distinct participant clusters (see Figure 1A and 1B). The *CMV seropositive senescent* cluster, highlights an often-recurring and frequently discussed result: significantly older, cytomegalovirus (CMV) positive individuals show senescence traits for CD4+ and CD8+ effector memory T cells alike, along with exhaustion for CD4+ and CD8+ effector cells and loss of the naive T cell compartments (see Figure 1C and 1D), in our statistical testing (conservative Bonferroni adjusted p-values < 0.05). In contrast, the *Naive group* represents a segment of the population with more naive T cells. Significantly younger individuals were here characterized by the highest levels of naivety markers for T lymphocytes and activated NK (Natural Killer) cells, while presenting low levels of exhaustion and central memory T cells (see Figure 1C). A separate *Elderly group* (CMV IgG titers adjusted p-value < 0.05) instead draws a picture of an exhausted phenotype. Both exhausted central memory CD4+ and CD8+ T cell populations were overrepresented, coupled with increased levels of CD4+ regulatory T cells. Interestingly, the *Elderly group* exhibited a significant double-negative T cell (DNT) abundance, suggesting a correlation between DNT and T cell exhaustion in humans. *Monocyte-driven* individuals showed an increase monocyte-specific markers (CD11cint) along with significant CD4+ specific signatures, namely CD4+ naive T cells and activated CD4+ T effect cells, while presenting lower levels of exhausted CD4+ of Th2 differentiated cells, CD4+ T regs and LAG3+ (Lymphocyte Activation Gene 3) B cells. The *FAS cluster* represented individuals with high abundant Fas+tim3+ monocytes and significantly low NK cell numbers, naive T cell numbers and DNT counts. Finally, the smaller *LAG3 cluster (only four i*ndividuals) positioned itself at the other end of the spectrum compared to the *Monocyte-driven* group, with reversed significant patterns.

**Figure 1:**
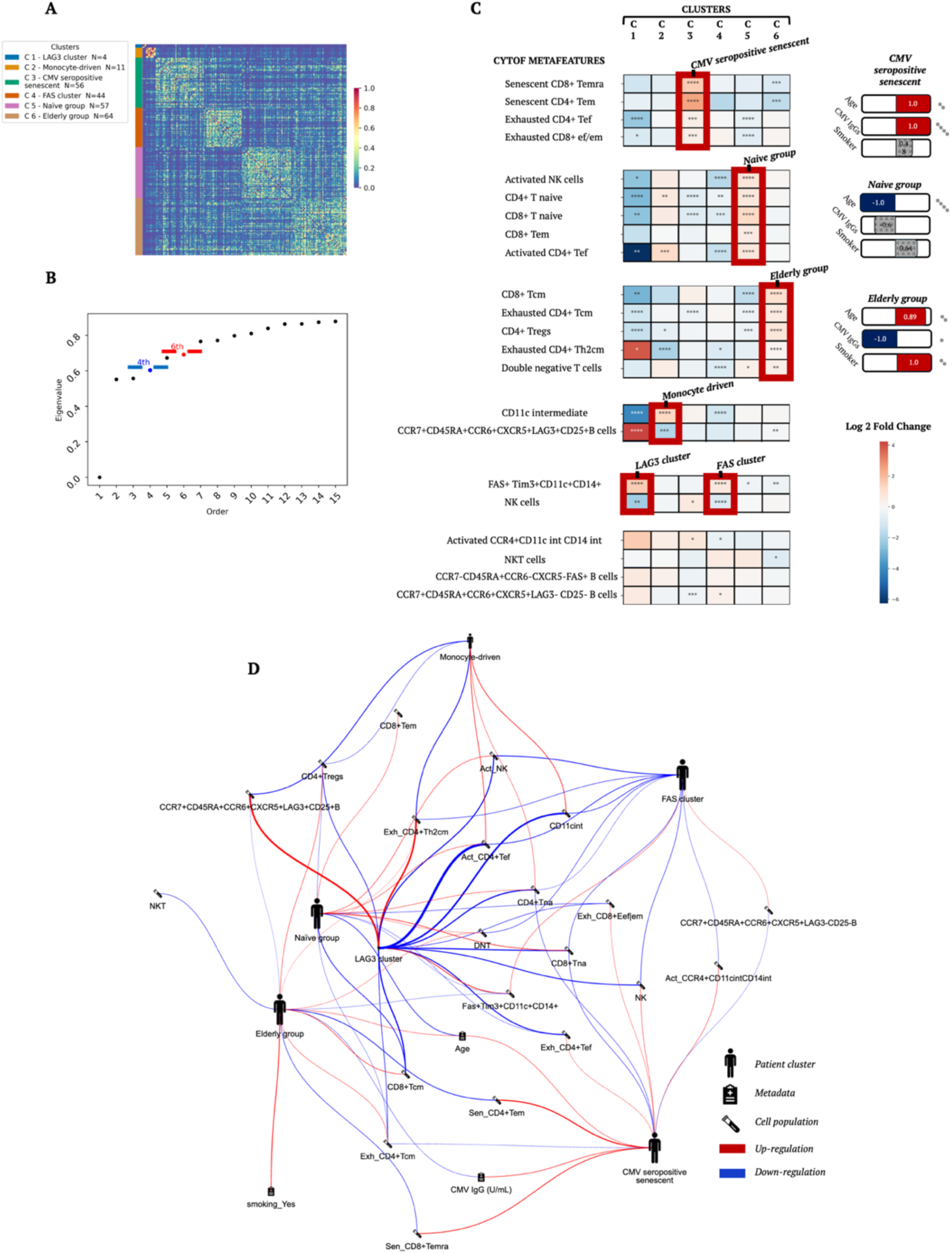
CyTOF profiles grant a robust starting point for multimodal analyses. (A) Squared Euclidean distance matrix for the CyTOF modality, with distances minmax-rescaled between 1 and 0 for visualization purposes. (B) The top 15 smallest eigenvalues of the Laplacian matrix, sorted in ascending order, highlight the connectivity profile of the underlying data. Two “jumps” are present, after the fourth and sixth eigenvalues, as candidate cutoff points for the optimal number of clusters. (C) Heatmap of the statistical testing results contrasting 22 cytometry patterns (metafeatures) for each patient cluster against the remainder of the cohort (total of n = 236). The hue indicates, for each feature, the log2 ratio of its mean expression in the cluster over the mean expression in the background. P-values were obtained from Mann-Whitney u tests assessing significance adjusted for multiple testing with the Bonferroni method. Normalized coefficients of a logistic regression on demographic features and their significance are also reported in bar plots for clusters for which at least one significant coefficient is present. Non-significant features are grayed-out. Asterisks indicate statistical significance (∗p < 0.05, ∗∗p < 0.01, ∗∗∗p < 0.001, and ∗∗∗∗ p < 0.0001) and non-significance when missing. Abbreviations: Temra, terminal effector memory T cells; Tem, effector memory T cells; Tef, effector memory T cells; Tcm, central memory T cells; NK, natural killer cells; NKT, natural killer T cells; int, intermediate.

### Gut microbiome screening highlights enterotypes characterized by BMI profiles and community diversity

Next, we performed 16s rRNA sequencing on fecal material from 182 individuals (Figure S1, Table S1), yielding in total 7954 observed amplicon sequence variants (ASVs). After removal of ASVs present in less than 10% of the cohort, the final selection included 387 distinct ASVs at the genus level.

Separate Logistic regression (LR) analysis of the gut microbiome modality led to the identification of BMI as the main metadata feature associated with cluster membership (only clinical data feature reporting coefficient significance, adj. P-value < 0.05, Figure 2A). Four microbiome clusters were identified with a stark difference in bacterial richness and evenness profiles among them (Shannon entropy metric, see Figure 2B). Differences in the microbial community composition between the identified clusters was confirmed by a significant (p-value < 0.0001) non-parametric multivariate statistical permutation test (PERMANOVA) with R2 of 0.126 (Figure 2C). The investigation of shared enterotypes was addressed with Gene Set Enrichment Analysis (GSEA)^15^, by examining enrichment on a bacterial family level.

**Figure 2:**
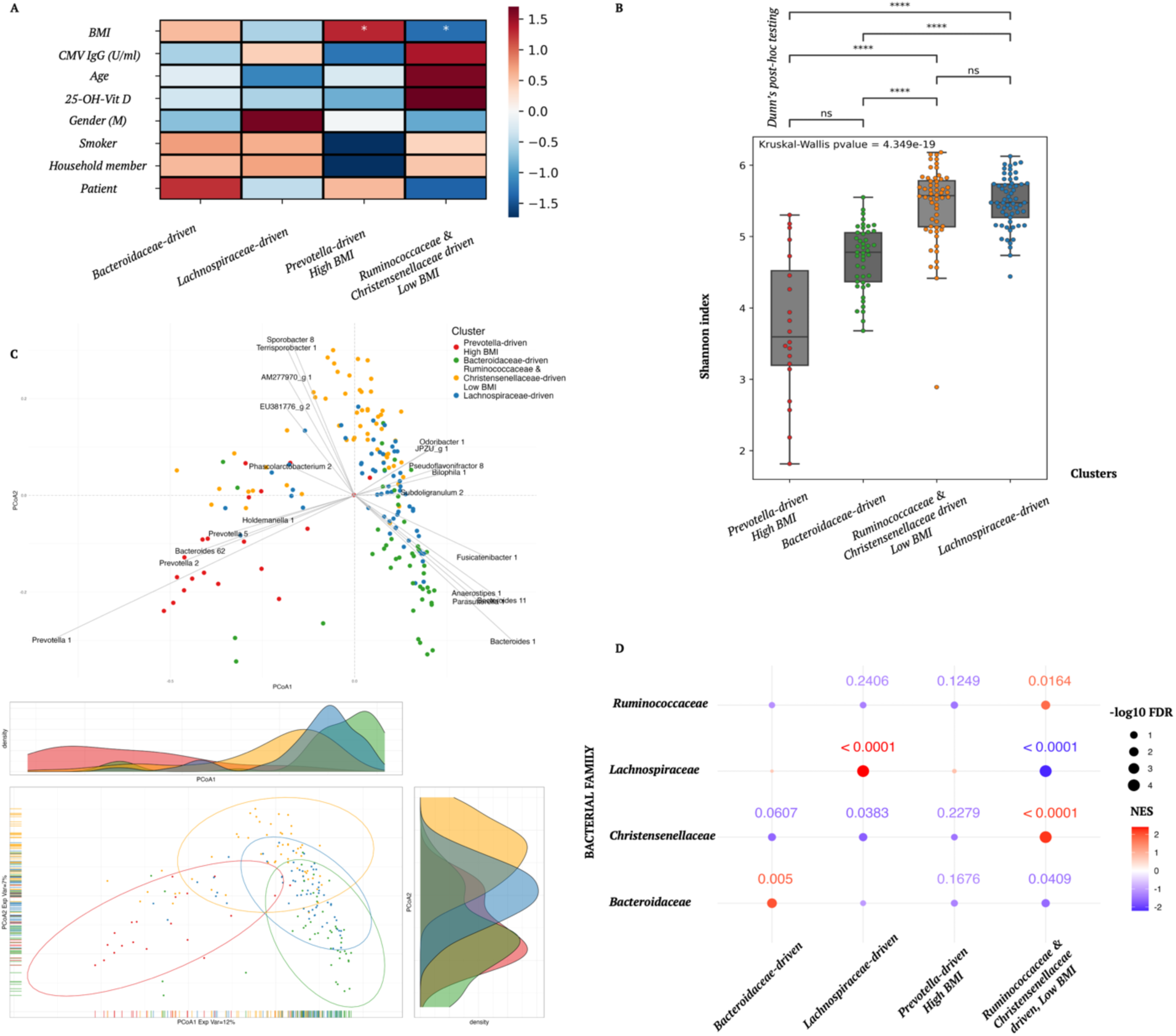
Gut microbiome screening highlights enterotypes characterized by different BMI profiles. (A) Results of a logistic regression (LR) on the metadata for the Microbial unimodal case (n = 182). The hue indicates the LR coefficients obtained by comparing each cluster against the background cohort. Statistical significance of the coefficients is indicated with asterisks (*p < 0.05, missing = not significant). (B) Shannon index distribution boxplot is shown for each cluster identified. Box plot center lines represent the median values; lower and upper hinges correspond to the first and third quartiles, lower and upper whiskers extend to the smallest and largest value respectively, reaching at maximum 1.5 times the interquartile range. Kruskal-Wallis test’s p-value is reported on the plot while Bonferroni corrected Dunn’s post hoc tests statistical significance is indicated with asterisks (ns = not significant, *p < 0.05, ∗∗p < 0.01, ∗∗∗p < 0.001, and ∗∗∗∗ p < 0.0001). Principal Coordinate Analysis (PCoA) performed with Bray-Curtis distance shows stark separation of the enterotypes in the low dimensional space analysis, indicating significant intra-cluster differences in bacterial diversity. (C) (top) For each quadrant, the top five most strongly correlated ASVs with the latent dimensions (PCoA1 and PCoA2) are shown. The two legs on the x and the y axes, identified by the corresponding ASV (amplicon sequence variants) gray line, represent the correlation of the bacterial abundance with PCoA1 and PCoA2 respectively. (bottom) Density plots for each axis and ellipses representing the 95% confidence interval assuming a multivariate t-distribution are shown. PCoA1 captured 12% of the explained variance while PCoA2 captured 7% explained variance. (D) Microbial family-wise enrichment results in a one-vs-rest setup. Results of a Gene Set Enrichment Analysis (GSEA) performed on the bacterial families are reported. Only results from families of a large enough size for GSEA to not reject a priori the set are shown. The color scale represents the normalized enrichment score, the dot size is the –log 10 False Discovery Rate (FDR). If the FDR fell below the standard GSEA threshold of 0.25 its value is shown next to the respective dot, rounded up to the fourth decimal place.

A *Prevotella-driven* cluster showed the most elevated BMI (LR coefficient = 0.67 with Bonferroni corrected p-value = 0.028, Figure 2B) and the lowest median entropy (*Kruskal-Wallis, p < 0.05,* Figure 2A). The cause of low diversity in this cluster was traced to a few *Prevotella* ASVs representing up to 70% of the gut bacterial flora of these individuals. However, in our dataset, this *Prevotella* enterotype seemed to be only a minor fraction of the global participant population (circa 10% - 20% individuals, see Figure S2A). A second cluster showed the opposite relation between BMI and median entropy coupled with a significant increase in family-wise expression of *Ruminococcaceae* and *Christensenellaceae bacteria* (NES > 0 and FDR < 0.25, Figure 2D). The lower BMI, high bacterial diversity cluster denoted as the *Ruminococcaceae-and-Christensenellaceae-driven cluster* was here hypothesized to be a homeostatic group that can be seen as the core of the quiescent fraction of the cohort in the following analyses.

On the other hand, a *Bacteroidaceae-driven* and a *Lachnospiraceae-driven* cluster occupied intermediate conditions in terms of BMI and bacterial diversity, with the *Bacteroidaceae-driven* cluster having lower entropy among the two.

### Blood immune cell repertoire and gut microbiome show little overlap in their clustering of participants

Next, we investigated how the CyTOF and gut microbiome unimodal analyses were overlapping to identify a consensus signature. A cross comparison of the repartition into groups for the 101 individuals of which both modalities were available was thus drawn (Figure S1, Table S1), revealing no transversal relationship between the two distinct clustering outcomes (see Figure 3A and 3B). The observed Adjusted Rand Index between the two unimodal analyses labels was 0.0163 (p = 0.1309, permutation test). This result suggests that these two types of data exert distinct influences on the clustering process, producing minimal concordance. Moreover, beyond the p-value originating from the permutation test being not significant at the usual 0.05 value, the overall distribution falls around the zero value, confirming the negligeable overlap between the CyTOF and microbiome unimodal labeling.

**Figure 3:**
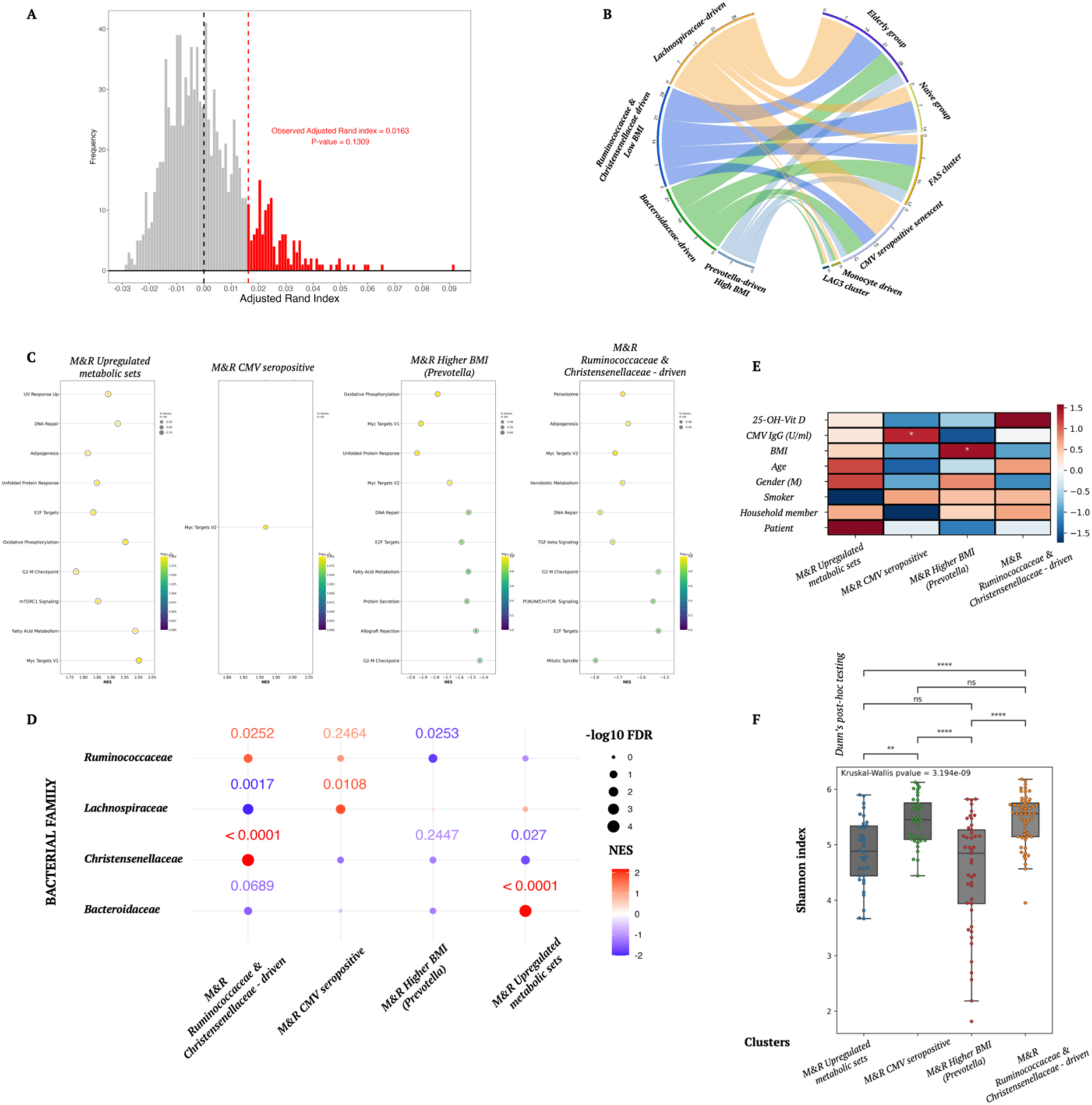
CyTOF and gut microbiome show little overlap in their clustering, however, the transcriptome bridges the gap between the two modalities. (A) Results of a permutation test conducted on the Adjusted Rand Index (ARI) between the CyTOF unimodal clustering and the Microbiome unimodal clustering (n = 101). The observed ARI was first calculated on the unmodified two sets of labels. The labels were then permuted one thousand times, and the ARI was recalculated at each step producing the distribution shown. A p-value was then calculated as the fraction of permutations that produced an ARI greater than the observed value (red hue). The dashed red line shows the observed ARI, while the dashed black line represents the zero ARI. (B) Chord diagram depicting the Microbiome (left) and CyTOF (right) repartitions of the respective unimodal clusters. Colored flows based on the CyTOF labels show the overlap between the two modalities. (C) Gene Set Enrichment Analysis (GSEA) Normalized Enrichment Scores (NES) in a one-vs-rest setup for the bimodal Microbiome & RNAseq case (n=171). Only gene sets enriched at a False Discovery Rate (FDR) < 0.25 are here reported. The hue represents the FDR while the dot size shows the ratio of genes in the gene set after filtering out those genes not in the expression dataset. (D) Microbial family-wise enrichment results in a one-vs-rest setup. Results of a GSEA performed on the bacterial families are reported. Only results from families of a large enough size for GSEA to not reject a priori the set are shown. The color scale represents the normalized enrichment score, the dot size is the –log 10 FDR. If the FDR fell below the standard GSEA threshold of 0.25 its value is shown next to the respective dot, rounded up to the fourth decimal place. (E) Results of a logistic regression (LR) on the metadata for the bimodal Microbiome & RNAseq case. The hue indicates the LR coefficients obtained by comparing each cluster against the background cohort. Statistical significance of the coefficients is indicated with asterisks (*p < 0.05, missing = not significant). (F) A boxplot is shown for each cluster identified. Box plot center lines represent the median values; lower and upper hinges correspond to the first and third quartiles, lower and upper whiskers extend to the smallest and largest value respectively, reaching at maximum 1.5 times the interquartile range. Kruskal-Wallis test’s p-value is reported on the plot while Bonferroni corrected Dunn’s post hoc tests statistical significance is indicated with asterisks (ns = not significant, *p < 0.05, ∗∗p < 0.01, ∗∗∗p < 0.001, and ∗∗∗∗ p < 0.0001).

### The blood transcriptome captures orthogonal patterns from the gut microbiome and immune cell composition

Blood transcriptomics data was also collected for a total of 317 individuals (Figure S1, Table S1). We recalculated patient similarities, combining the RNAseq modality in turns with the other data types in a Similarity Network Fusion (SNF) calculation, exploring the added value of the RNAseq dimension to substantiate the microbiome-host immune cells cross talk and illuminate the proinflammatory link between different aspects of the host microenvironment.

The 171 individuals available for the Microbiome – RNAseq integration (M&R) (Figure S1, Table S1) identified four clusters (Figure 4). The *M&R Ruminococcaceae and Christensenellaceae* cluster was associated with downregulation of pathways such as adipogenesis, xenobiotic metabolism, mTOR signaling, Myc targets, DNA repair, peroxisome (NES < 0 and FDR < 0.25, Figure 3C). This downregulation paired with upregulation of the Ruminococcaceae and Christensenellaceae families (NES > 0 and FDR < 0.25, Figure 3D) supported the quiescent phenotype. A highly metabolic group, denoted as the *M&R upregulated metabolic* cluster, can be identified as the progression of the unimodal *Bacteroidaceae-driven* cluster with additional fatty acid metabolism, adipogenesis and oxidative phosphorylation pathway upregulation. Patients from the previous *Prevotella-driven* cluster (rich in *Prevotella 1 and Prevotella 2*) prevalence were incorporated in the *M&R higher BMI* cluster, due to a significant BMI increase over the remaining individuals (Figure 3E). The inclusion of RNA in the analysis, however, meant that more individuals were grouped in this cluster, hence reducing the statistical significance of the community entropy difference (no significance in the comparison against the *M&R upregulated-metabolic* cluster – Figure 3F). The GSEA transcriptomic enrichment scores for the *M&R higher-BMI* cluster surprisingly elucidated an overall downregulation for many metabolic gene sets in contrast to the *M&R upregulated-metabolic* cluster. Finally, the *M&R CMV-seropositive* cluster, characterized by an overabundance of CMV positive individuals, is interestingly the only group for which the *Lachnospiraceae* family is overexpressed (in conjunction with *Ruminococcaceaae*)

**Figure 4:**
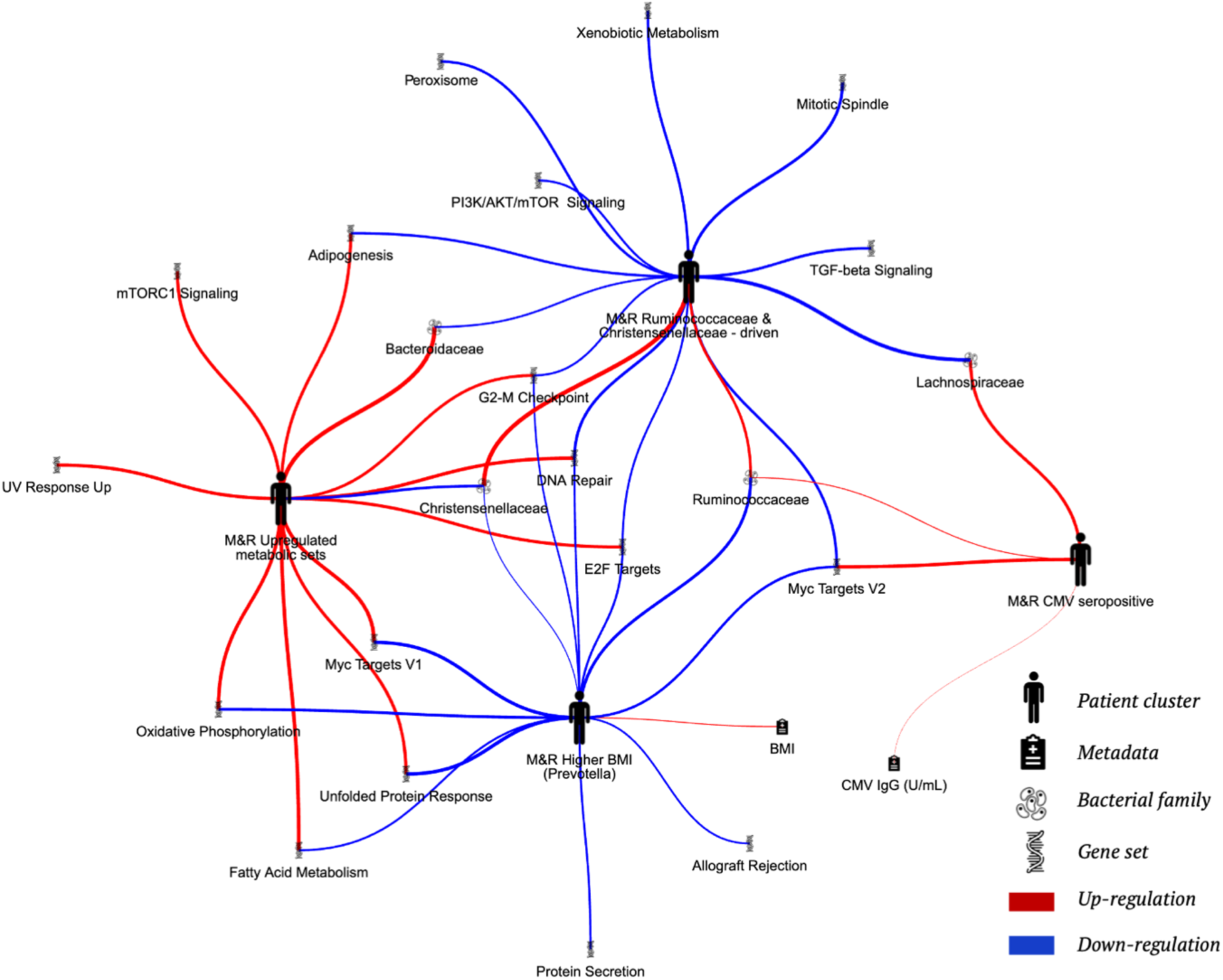
The transcriptome bridges the gap between CyTOF and gut microbiome. Spider plot drawing a summary of the bimodal integration of RNAseq and Microbiome (n = 171). Only significant results for the respective tests are reported. Red and blue arches respectively indicate significantly positive and negative regulation of the corresponding variable in the cluster. Icons representing patient clusters are scaled with respect to cluster size. The arch width has been scaled per analysis and indicates the effect size. Abbreviations: M&R, Microbiome and RNAseq.

A similar clustering analysis can be performed at the level of CyTOF - RNA integration (C&R) (Figure 5), based on 177 individuals (Figure S1, Table S1). This resulted in the bimodal cluster C&R 4 derived from the CyTOF unimodal *CMV seropositive senescent* group (Figure 1). The RNA layer solidified this result through the positive expression of many inflammatory gene sets, including upregulation of interferon alpha and gamma responses, potentially indicative of an ongoing anti-viral response (Figure S3A). This finding indicates that CMV-seropositivity might, at least in this cluster, not be dormant, but rather correlated with active CMV presence. Additionally, C&R 4 displayed further upregulation of UPR, OxP and mTOR signaling.

**Figure 5.**
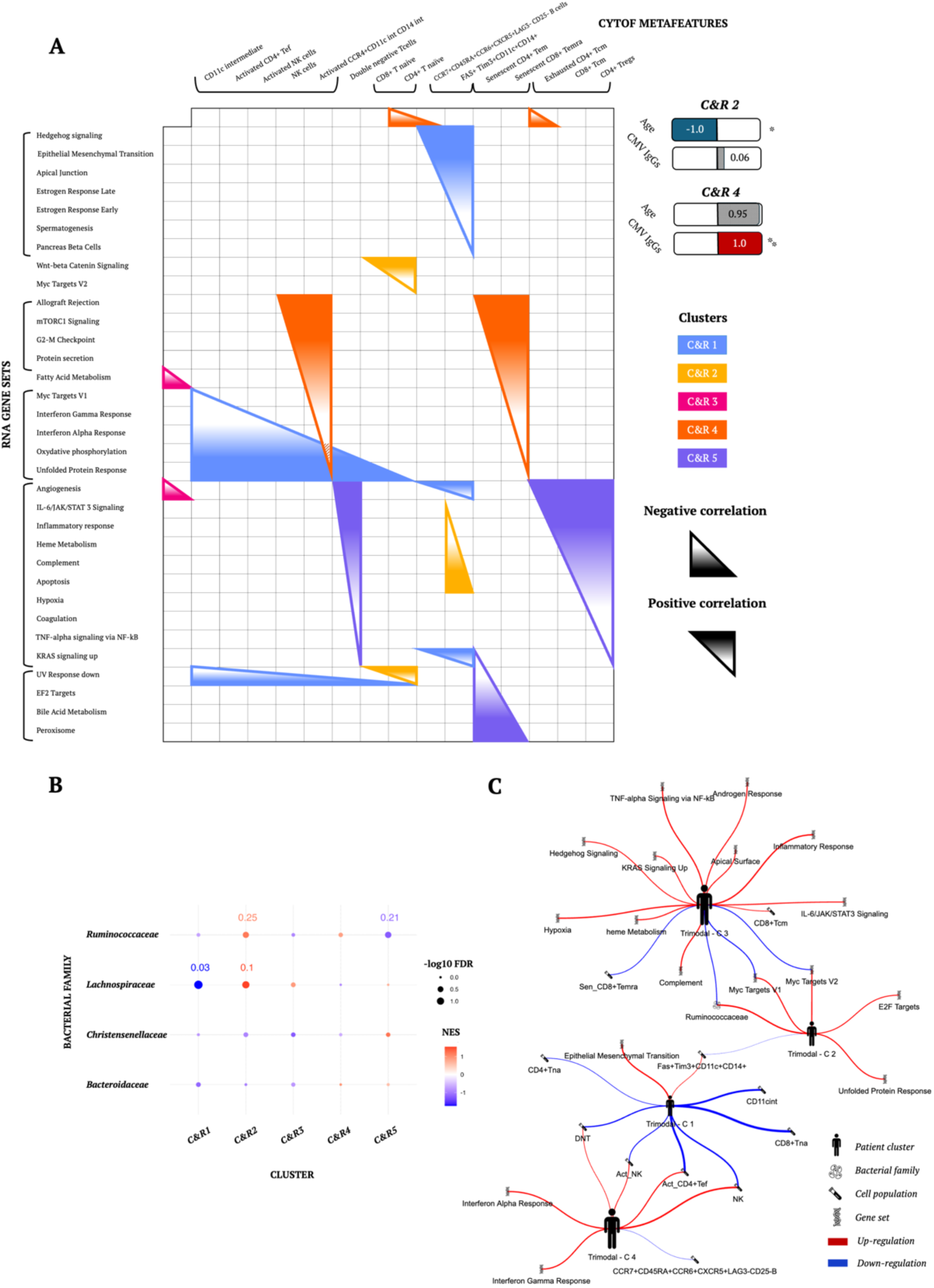
The blood transcriptome captures orthogonal patterns from the gut microbiome and immune cell composition. (A) Summary of the bimodal integration of RNAseq and CyTOF (n=177). A triangle is drawn to highlight features from the two datasets that are negatively or positively co-expressed. The orientation and the gradient of the triangle indicate the sign of the relationship. Only significant results for the respective tests are reported. Normalized coefficients of a logistic regression performed on the metadata are also reported when significant for at least one patient cluster. Statistical significance is indicated with asterisks (*p < 0.05, ∗∗p < 0.01, ∗∗∗p < 0.001, and ∗∗∗∗ p < 0.0001). (B) Ruminococcaceae enrichment correlates with systemic wellness by the additional overlay of the gut microbial component, which was however not used in the clustering computation. Results of a Gene Set Enrichment Analysis (GSEA) performed on the bacterial families are reported. Only results from families of a large enough size for GSEA to not reject a priori the set are shown. Namely: Ruminococcaceae (116 ASVs), Lachnospiraceae (104 ASVs), Christensenellaceae (29 ASVs) and Bacteroidaceae (25 ASVs). The color scale represents the normalized enrichment score, the dot size is the –log 10 False Discovery Rate (FDR). If the FDR fell below the standard GSEA threshold of 0.25 its value is shown next to the respective dot, rounded up to the second decimal place. (C) Spider plot drawing a summary of depicts the trimodal integration of RNAseq, Microbiome and CyTOF (n=97). Only significant results for the respective tests are reported. Red and blue arches respectively indicate significantly positive and negative regulation of the corresponding variable in the cluster. Icons representing patient clusters are scaled with respect to cluster size. The arch width has been scaled per analysis and indicates the effect size. Abbreviations: Temra, terminal effector memory T cells; Tem, effector memory T cells; Tef, effector memory T cells; Tcm, central memory T cells; NK, natural killer cells; NKT, natural killer T cells; DNT, double negative T cells; int, intermediate; Act, activated; na, naive; C&R, CyTOF and RNAseq.

In contrast, C&R 2 includes younger individuals with more naive T cells (both CD4 and CD8) and less Fas+Tim3+ monocytes (Figure S3B and S3C) and presenting downregulation in metabolic and inflammatory pathways (inflammatory response, apoptosis, heme metabolism, IL-6) but, strikingly, an upregulation of Wnt-beta catenin signaling. This last pathway has been identified to play a critical role in immune-related disorders^21^. Furthermore, CD14+ cells and coincident JAK/STAT3 pathway enrichments have been shown to be a recurring ’M1-like’ hyper inflammatory, hyper metabolic phenotype^22^. Here both were downregulated.

C&R1, distinguished by seemingly migratory cells, which are marked by the expression of three cytokine receptors (CCR7, CCR6, CXCR5), also interestingly showed significance for Epithelial Mesenchymal transition, which corroborates the hypothesis of enhanced cellular migration in some individuals^23^.

When overlaying the microbial component, without using this modality for similarity clustering, on the CyTOF and RNA bimodal integration, we detected in the homeostatic C&R 2 a significant positive NES for the *Ruminococcaceae* family (Figure 5B), once again corroborating the conclusion on the positive association of this bacterial family on health prospects^24^. The opposite enrichment could be found in C&R 5. Lastly, a cluster of 29 individuals emerged (C&R 3), characterized exclusively by reduced expression of *Angiogenesis* and *Fatty acid metabolism genes*.

When combining directly all three modalities in the calculation to obtain tri-modal clusters (n = 97, Figure S1, Table S1), many patterns were preserved but were accompanied by a loss of statistical power (Figure 5C). *Trimodal C2*, being the younger, *Ruminococcaceae* upregulated group, featured the highest median bacterial entropy, as well as the lowest BMI (not significant). This group can be seen as the most consistent pattern across integrations (Figure S4A) and is maintained no matter the strategy chosen. *Trimodal C3* showed a significant absence of the *Ruminococcaceae* family while most inflammatory gene sets were upregulated, combined with cytotoxic CD8+ Tcm cells (Figure 5C) and a higher BMI (although not significant – Figure S4B). The key factor that permeated the RNA-Microbiome integration, BMI, was expectedly lost due to the cluster consistency being disrupted by the averaging with the CyTOF layer.

This loss of consistency is replicated by the disappearance of a significant CMV effect in shaping the immune response that previously characterized the CyTOF unimodal analysis and the integration of CyTOF and RNA. *Trimodal C4* represented the immuno-activated cluster in this tripartite integration, from the RNAseq perspective with anti-viral interferons alpha and gamma and from the CyTOF dimension with NK and activated NK cells, activated CD4+ effector cells and DNT cells (Figure 5C). However, CMV and senescence markers were both elevated but not significant (Figure S4B and S4C).

### TCR repertoires show skewing towards putative SARS-CoV-2 clusters

Key aspects of the human adaptive immunity compartment can only be harnessed by delving deeper into the statistical properties of the T cell receptor (TCR) repertoires. TCR repertoires collected from blood-derived bulk CD4+ and CD8+ T cells from 99 individuals allowed us to explore this aspect (Figure S1, Table S1). TCR-based patient clusters built on the co-expression of similar CDR3 amino acid sequences, appeared to correlate with SARS-CoV-2-specific traits (Figure 6A, see Methods). Members of the cohort were indeed segregated into four well-distinguished groups, with differences in frequency and specificity matching for SARS-CoV-2epitopes. However, modules constructed in the current manner could be affected by an intrinsic bias.

**Figure 6:**
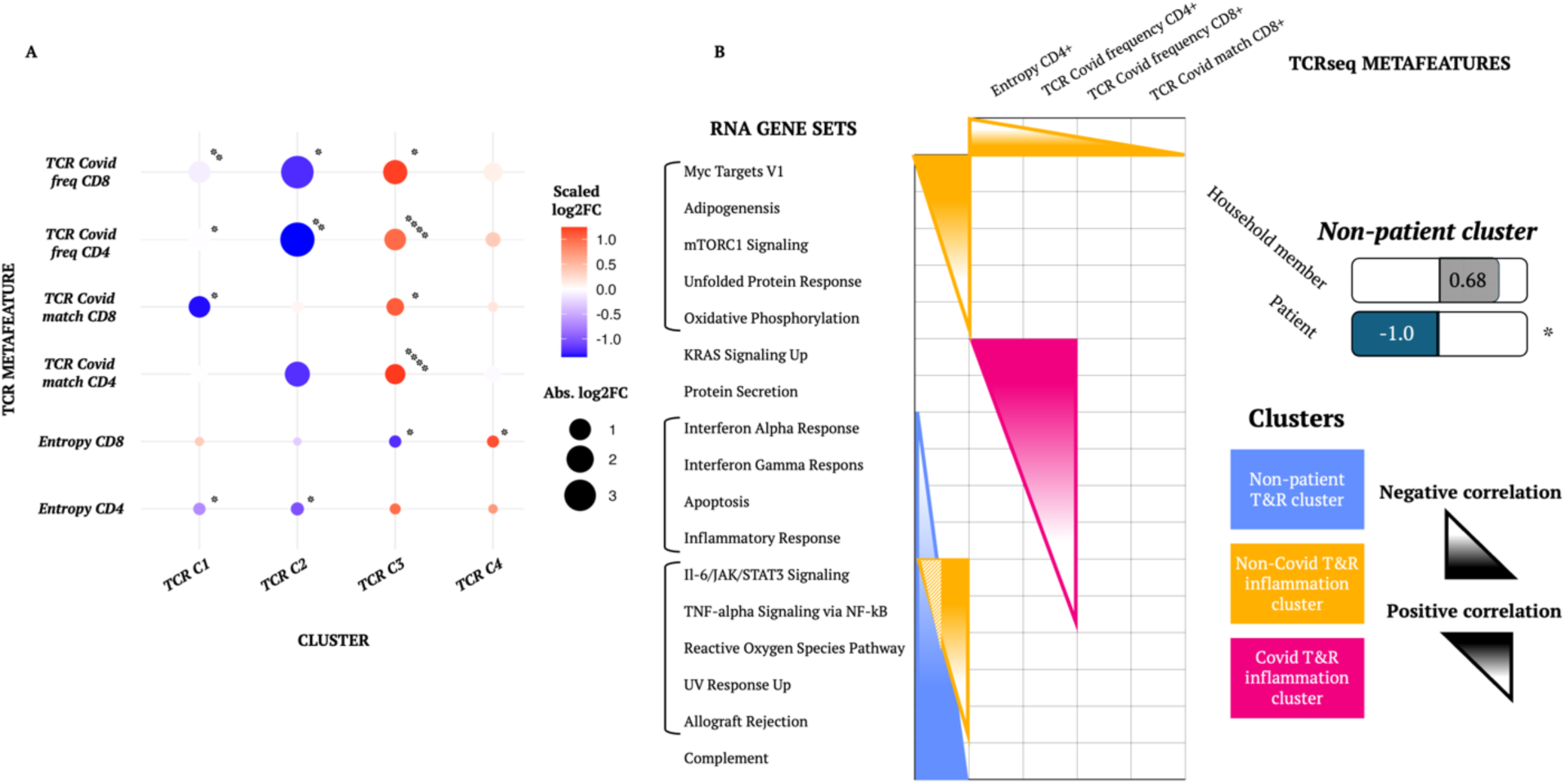
TCR repertoires show skewing towards putative SARS-CoV-2 clusters. (A) Results on the unimodal TCRseq layer for each TCR-specific statistic (metafeature). The hue indicates the log2 ratio of its mean expression in the cluster over the mean expression in the background. Values were scaled between –1 and +1 for visualization purposes (hue) while the size of the dots represent the absolute log2 fold change value. P-values were obtained from Mann-Whitney u tests assessing significance adjusted for multiple testing with the Bonferroni method. (B) Summary of the bimodal integration of RNAseq and TCRseq (n=86). A triangle is drawn to highlight features from the two datasets that are negatively or positively co-expressed. The orientation and the gradient of the triangle indicate the sign of the relationship. Only significant results for the respective tests are reported. Normalized coefficients of a logistic regression performed on the metadata are also reported when significant for at least one patient cluster. Asterisks indicate statistical significance (∗p < 0.05, ∗∗p < 0.01, ∗∗∗p < 0.001, and ∗∗∗∗ p < 0.0001) and non-significance when missing. Abbreviations: T&R, TCRseq and RNAseq.

### Data integration across all layers is driven by inflammation markers

Regardless of the origin, the main recurring driver of heterogeneity in the cohort must be sought in the inflammatory phenotype. We used again the blood transcriptome dimension to add depth and interpretability, in this instance to the TCR modules (n = 86, Figure S1, Table S1). Integrating the RNA and TCR component (T&R) revealed that SARS-CoV-2 convalescent adults are, even more than 3 months after a SARS-CoV-2 infection, still in an inflammatory state (both IFN and non-IFN based). In this analysis, the *Inflammation T&R cluster* likely encompassed individuals experiencing a sustained inflammatory state caused by the long-term Covid-19 aftermath or persistence of SARS-CoV-2-specific epitopes in their system (Figure 6B).

A second inflammatory cluster was present (*Non-Covid T&R inflammation cluster*), although clearly not related to Covid19 due to the negative associations of frequency for SARS-CoV-2-specific CD4+ TCRs, the low respective entropy but also significantly low TCR SARS-CoV-2 matching and frequency of SARS-CoV-2-specific CD8+ T cells (Figure 7A and 7B). The *Non-Covid T&R inflammation cluster* was also remarkably not IFN-driven, in contrast to the previous *Covid T&R inflammation cluster*. The *Non-patient T&R cluster* involved mostly those individuals that are not classified as “SARS-CoV-2 patients” thus this group is mostly made up of healthy, non-infected, non-inflammatory, individuals. Only 35% of the *Non-patient T&R cluster* have contracted SARS-CoV-2 compared to 76% of the remainder (Fisher exact test p-value = 0.001). However, for one group of 20 individuals, denoted as *T&R cluster 3*, no significant associations were found (Figure *7C*).

**Figure 7:**
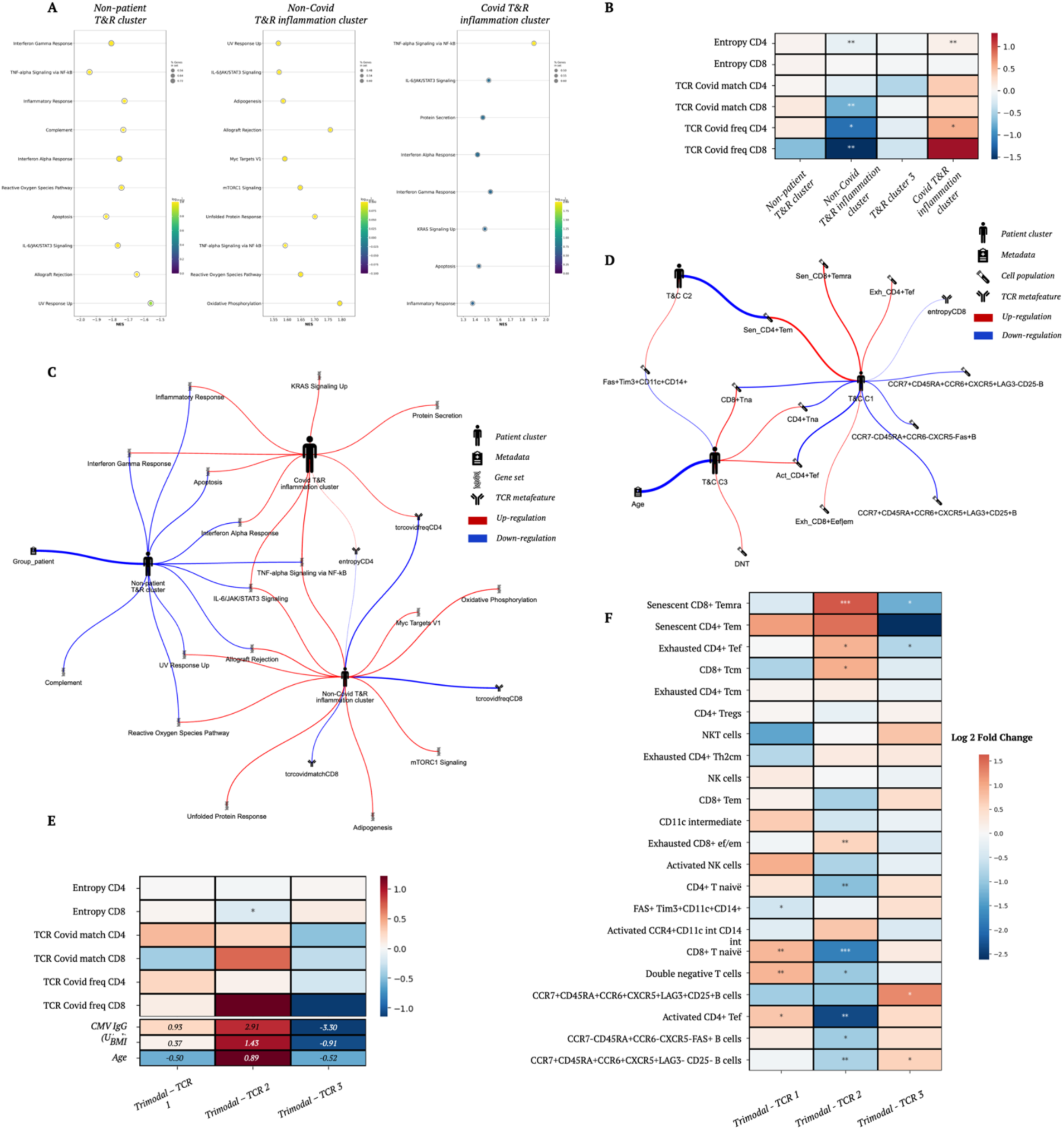
Data integration across all layers is driven by inflammation markers. TCRseq and RNAseq bimodal integration (n=86). (A) Gene Set Enrichment Analysis (GSEA) Normalized Enrichment Scores (NES) results. Only gene sets enriched at a False Discovery Rate (FDR) < 0.25 are here reported. The hue represents the FDR while the dot size shows the ratio of genes in the gene set after filtering out those genes not in the expression dataset. (B) Results for the enrichment testing of TCR-specific statistic (metafeature). The hue indicates the log2 ratio of its mean expression in the cluster over the mean expression in the background. The hue represents the log2 fold change value. P-values were obtained from Mann-Whitney u tests assessing significance adjusted for multiple testing with the Bonferroni method. (C) Spider plot drawing a summary of the bimodal integration of RNAseq and TCRseq (n=86). Only significant results for the respective tests are reported. Red and blue arches respectively indicate significantly positive and negative regulation of the corresponding variable in the cluster. Icons representing patient clusters are scaled with respect to cluster size. The arch width has been scaled per analysis and indicates the effect size. (D) Spider plot drawing a summary of depicts the bimodal integration of TCRseq and CyTOF (n=47). Only significant results for the respective tests are reported. Red and blue arches respectively indicate significantly positive and negative regulation of the corresponding variable in the cluster. Icons representing patient clusters are scaled with respect to cluster size. The arch width has been scaled per analysis and indicates the effect size. Trimodal integration of RNAseq, CyTOF and TCRseq data (n=45). (E) (top) Results for each TCR-specific statistic (metafeature). The hue indicates the log2 ratio of its mean expression in the cluster over the mean expression in the background. P-values were obtained from Mann-Whitney u tests assessing significance adjusted for multiple testing with the Bonferroni method. (bottom) Results of logistic regression on the metadata for the trimodal clusters. Values of the coefficients are reported, standard significance (0.05) after correction was not reached by any feature. (F) Heatmap of the statistical testing results contrasting 22 cytometry patterns (metafeatures) for each patient cluster against the remainder of the cohort. The hue indicates, for each feature, the log2 ratio of its mean expression in the cluster over the mean expression in the background. P-values were obtained from Mann-Whitney u tests assessing significance adjusted for multiple testing with the Bonferroni method. Asterisks indicate statistical significance (∗p < 0.05, ∗∗p < 0.01, ∗∗∗p < 0.001, and ∗∗∗∗ p < 0.0001) and non-significance when missing. Abbreviations: Temra, terminal effector memory T cells; Tem, effector memory T cells; Tef, effector memory T cells; Tcm, central memory T cells; NK, natural killer cells; NKT, natural killer T cells; int, intermediate; exhausted; Sen, senescent; na, naive; DNT, double negative T cells; T&C, TCRseq and CyTOF; T&R, TCRseq and RNAseq.

Next, we fused the TCR layer similarities with the blood immune cell repertoire data layer (via CyTOF) to investigate potential correlates of SARS-CoV-2 specificity with the host’s distribution of immune cell populations (TCR and CyTOF integration – T&C; n = 47). Here T&C C3, strongly recaptured the pattern represented by younger individuals who showed naive T cell markers in the CyTOF unimodal analysis (the *Naive group* of Figure 1). T&C C3 did indeed hold CD4+T naive, CD8+ T naive, DNT and activated CD4+ T effector cells overrepresentation (Figure *7*D). Interestingly, individuals in T&C C1 showed the lowest CD8+ entropy. From a CyTOF point of view, they were confirmed to be part of the highly enhanced senescent cluster with less naïve T cells. Overall, this can be interpreted as a senescence association with CD8+ TCR repertoire narrowing, which is in line with the reduced sequence diversity in the SARS-CoV-2 specific T cell response and can be considered its natural explanation.

By expanding the previous analysis with the RNA-seq expression data (n = 45, Figure S1, Table S1), thus producing a trimodal integration of RNAseq, CyTOF and TCRseq data, we largely confirmed the previous findings while revealing a lowly inflamed, healthier phenotype in the *Trimodal TCR 3* cluster (Figure S5). The lowest CD8+ entropy marker in *Trimodal TCR 2* cluster was maintained while also presenting the highest CD8+ TCR Covid matching and frequency, highest age, BMI and CMV IgG titer, albeit most of these features were not show significant differences (Figure *7*E). For *Trimodal TCR 3* we must highlight the recurrent anticorrelated relationship between senescence/exhaustion and two novel B cell populations, also present in T&C C1 (*Figure 7F*). For this group, the downregulation of several inflammatory pathways, mainly INF alpha and gamma and IL-6, supported the low terminal differentiation of the adaptive compartment.

## Discussion

### Complexities in multimodal data dynamics lead to divergent perspectives

In this multimodal study where we profiled 394 individuals, we found evidence of how different data type modalities give information on complementary/orthogonal aspects of the status of the individual. In isolation, they disagree on the distinction between homeostatic and dysbiosis conditions. This is clearly shown by the lack of overlap between the blood immune cell repertoire (measured via CyTOF) and fecal Microbiome unimodal analyses. This illustrates the large challenge in identifying which profile would be most desirable for one’s health prospects.

Features that, in isolation, are seemingly conducive to a healthy phenotype, when accompanied by complementary information, instead lead to a more fine-grained fragmentation of the population, possibly suggesting continuity. This finding aligns with what has been found in a prior study^1^, and thus the lack of reliable metrics for assessing immune system health still stands as a major challenge. Prior analysis of immune cell populations revealed a continuous distribution of individuals rather than discrete clusters, proposing that the composition of immune systems varies substantially between individuals but largely remains stable over time due to compensation and redundancy of its parts^16^.

Our observation is also described by a well-established phenomenon in statistics, wherein the effects of explanatory variables can change dramatically from a simple regression to a complex multiple regression. Instead, we leveraged the blood transcriptome to identify and interpret common drivers across the orthogonal modalities. Here, inflammation emerged as the recurring and most stable pattern that informed the separation of individuals.

In this framework it is crucial to employ holistic methods that can capture the complexity of the immunotype at once, recapitulating its branches in a unified systemic approach, given the assumption that immune mechanisms impact all data types.

### Ruminococcaceae enrichment correlates with systemic wellness

*Bacteroides* and *Prevotella* are known to be anticorrelated and antagonistic bacteria and this pattern was replicated in our fecal Microbiome unimodal taxonomic profile^25^. In particular, *Prevotella* has been reported to be a genus characteristic of healthy, plant-rich diets (high consumption of carbohydrates and fibers)^26^ but it has been also found in elevated numbers in inflammatory states and associated to obesity in some cases^27^, while the genus *Bacteroides* has been often reported in individuals consuming animal products-rich diets^28^.

Our results not only confirmed but also extended on the seminal Nature paper^2^, which introduced the concept of enterotype and that described the *Prevotella*, the *Bacteroides* and the *Ruminococcus-*based types. However, the topic of enterotypes has been greatly expanded with many more found in subsequent studies, also depending on the taxonomical level of analysis. Using a Principal Coordinate Analysis (PCoA) with Bray-Curtis distance as the metric of choice, we recapitulated the original authors’ projection, and we obtained a closely resembling distribution of the enterotypes. A significant PERMANOVA testing confirmed the statistical differences in bacterial abundances between the clusters.

Compared to the original study^2^ we also added a fourth enterotype, centered around the *Lachnospiraceae* family, and we linked membership to this enterotype to CMV seropositivity, possibly describing a novel axis between gut flora composition and CMV-carriage. Additionally, at all levels of the analysis, we found the *Ruminococcaceae* family to be linked with positive parameters such as younger age, lower BMI, an expansion in the CD8+ naive compartment and reduced inflammatory response, xenobiotic metabolisms and DNA damaging processes featuring as the most prominent. This is in line with reports that both the *Ruminococcaceae and Christensenellaceae* families show a positive influence on health^24,29^. Lastly, in line with the literature, BMI was identified as the main association to bacterial diversity throughout our analyses^30^.

### CyTOF robustness should guide immunological multimodal analysis

We initiated our analysis with a focus on the blood immune cell (CyTOF) modality due to its well-established nature, the manageable number of readily interpretable features generated and the overall satisfying compactness of its unimodal clusters in terms of silhouette scores and distinct profiles. The CyTOF toolset, being the best characterized among the techniques at our disposal, allows the information it provides to be extended to other biological levels. This provided a solid grounding for our study, allowing us to start the investigation with a strong foundation. Throughout our research, we consistently observed recurring findings derived from this modality. These persistent results underlined the significance and relevance of this data source in our data modelling. Complementing the CyTOF modality with gene expression, allowed us to recapitulate many markers related to T cell senescence that can be found in the literature such as oxidative stress, production of inflammatory cytokines (IFN - γ) and mTORC1 signaling to reduce autophagy^31^. We thus showed here that such results can be achieved through multi-view data integration on affordable bulk data.

We showed how CyTOF and Microbiome data have little to no overlapping behavior in the way they impact patient similarity. We thus needed to consider the blood immune cell repertoire and gut microbiome data as orthogonal datatypes. In the wake of our result, the wealth of papers indicating associations between the gut microbiota and blood immune cells and immunological disorders may seem to contradict our finding. Numerous are the papers supporting the idea of crosstalks between the gut microbiota and the immune system at large, making it possible to develop treatments tackling metabolic and immunological diseases by intervening on the gut flora^32^. The effects of beneficial intestinal bacteria have also been theorized to be contributing to systemic homeostasis, encompassing both the innate and the adaptive branches of the immune system, with dysbiosis possibly leading to autoimmunity^4^.

However, two aspects of the putative cross talks may be overlooked in the present investigation. Firstly, the longitudinal aspect is missing in our analysis and may be required to fully harness the interplay between bacteria and the immune system. Temporal changes in the microbial environment causing dysbiosis may affect more decisively the insurgence of metabolic diseases. Secondly, commensal and harmful bacteria have a deeper effect on the ecosystem in which they are embedded, directly shaping the intestinal mucosal immune system but less so the composition of immune cells in peripheral blood^33^. Nevertheless, it must be argued that our observation featuring the presented orthogonality is derived from pathogen-exposed humans instead of germ-free animal models that are often used to explore gut microbiota-immunity associations^34^.

### Cell types normally opaque to interpretation can be explained by way of multi-modal integration

The integration of the blood transcriptome with the blood immune cell repertoire (CyTOF) also allowed us to deconvolute cell populations with complicated kinetics such as Fas+Tim3+ monocytes. This cell type has been shown to present both a pro and an anti-inflammatory profile. Tim-3 has been proven to possess a regulatory effect on lymphocytes, as well as macrophages^35^, and, together with the Fas-mediated apoptosis of T cells^36^, they denoted an immuno-regulated pattern for a subset of participants from our cohort. In conjunction with our detected markers, activated CD4+ effector T cells as well as DNT and NK cells, were identified as being suppressed along with the transcriptomic interferon gamma response. On the other hand, in another subgroup, the absence of Tim-3 presenting cells determined an inflammation-deprived state in young individuals, thus illuminating the contrasting effects of Tim-3 as a marker of inflammation^37^. Fas+Tim3+ monocytes were recurrently detected as regulated in conjunction with a novel regulatory B cell subset (CCR7+CD45RA+CCR6+CXCR5+LAG3+CD24+ B cells)^38^. When both were downregulated, an overall “activated” cluster undergoing an underregulated immune response was determined. The nature of this downregulation could be driven by various causes including either low-grade disease or cancer predisposition, but at this stage it cannot be ascertained.

### Inflammation and viral pathogens leave a significant fingerprint on the hosts’ immune systems

The TCR modules strongly segregate individuals based on SARS-CoV-2 specific features. We can thus assume SARS-CoV-2 left a significant mark on the hosts’ immune systems by shaping the adaptive repertoire via persistent inflammation. However, shared HLA restriction could be the main driver of co-expression in the TCR modules, allowing for a biased clustering based on TCR-HLA association, as described by DeWitt et al^5^.

We also identified a cluster clearly not related to SARS-CoV-2 infection due to the negative associations of frequency for SARS-CoV-2-specific CD4 TCRs, the low respective entropy but also significantly low TCR SARS-CoV-2 matching and frequency of CD8 T cells. In this case, IFN gene sets were absent from the significant enrichment testing, confirming the hypothesis that it was not a viral-caused inflammation. TNF-alpha signaling via NF-kB and IL-6 gene sets were still upregulated but adipogenesis, allograft rejection, UV response, MYC targets were also present. Oxidative phosphorylation and Reactive oxygen species pathways as well as mTOR signaling suggest some sort of oxidative damage caused by, presumably, T cell-related metabolism [link].

CMV seropositivity is known to be a potent immunosenescence driver of the adaptive compartment^39–41^. We reported on a novel *Lachnospiraceae* association with a putative dormant CMV-carriage, and we repeatedly confirmed how a significant increase in serum CMV IgG levels, in conjunction with older age, were supported mainly by increased senescence and exhaustion of the CD4+ and CD8+ effector memory T cell subsets as well as systemic inflammation explained by the blood transcriptome.

### Study limitations

While the SNF methodology can be applied freely on different data, it is originally a kernel-based non-Bayesian network method, tailored for mRNA expression, DNA methylation and microRNA. SNF efficacy was demonstrated for these data layers but its performance on heterogenous data may be less consistent, lacking in generalizability. Even though we tried to reduce dimensionality issues to a minimum, this algorithm may inherently favor certain expression patterns and higher dimensional datasets, creating hard to detect biases.

Some additional drawbacks of the analysis derive from the use of spectral clustering. Firstly, the need to choose the number of clusters to be formed, a parameter not known a priori, may put this algorithm at a disadvantage when compared to methods that can more easily detect or discard outliers such as Density-Based Spatial Clustering of Applications with Noise (DBSCAN) and hierarchical clustering. Thus, spectral clustering assigns every point to a cluster, likely worsening the clustering result.

Furthermore, the limited number of samples in some integrations, compounded by the strict Bonferroni correction, may have lowered the power of association testing between metadata and cluster membership. The lack of more in-depth health parameters (i.e. current and past pathologies) and the dietary lifestyle may have also hidden confounding factors or part of the explanatory power, especially with regards to the microbiome layer.

## Methods

### Study cohort and datasets

A total of 569 individuals (Table S2), age range 15-85 (mean ± sd: 47.98 ± 15.68) were recruited and donated blood between September 2020 and July 2021. Metadata was collected as the only modality in the study reserved solely to perform regression analysis and not considered for patient similarity. The provenance of the individuals was documented as such: 254 controls (individuals without documented SARS-CoV-2 infection nor exposure), 164 covid-19 patients (participants who had recovered from COVID-19 (hospitalized or ambulatory) more than 3 months prior to enrollment and were screened prior to blood draws to make sure they were symptom-free and in a convalescent phase. On their visits, patients were asked to provide proof of positive testing for SARS-CoV-2, either via PCR or serological IgG testing and fever of 38°C or higher without other proven explanations) more than 3 months prior to recruitment), 151 household members (participants who lived in the same household with proven SARS-CoV-2 patients (who tested positive for SARS-CoV-2 by either PCR or IgG and had fever more than 3 months before). Household members did not have a confirmed SARS-CoV-2 infection, either by PCR or IgG.). Other features included: gender (321 female, 213 male), CMV IgG levels, Body-Mass Index (BMI) ranging from a minimum of 16.46 to a maximum of 53.35 (26 ± 4.77), Beck Depression Inventory score^42^ (BDI T score - 51.53 ± 9.4 with 434 missing values), Beck Anxiety Inventory score^43^ (BAI T score - 54.51 ± 11.65 with 421 missing values), binary variable indicating the occurrence of a previous Covid-19 infection for the individual if the information was available (297 negative, 208 positive), binary variable indicating if the patient experienced long Covid symptoms (110 negative, 89 positive), 25-hydroxy Vitamin D test results (25 OH Vit D - 21.27 ± 8.93 ng/mL) and a binary variable indicating the current smoker status (55 smokers, 478 non-smokers).

BDI and BAI values were the result of a standard Pearson assessment and are thus T scores based on the norm for non-clinical individuals (mean ± standard deviation: 50 ± 10) and not the usual Likert scale values. Occasional smokers and alternatives to cigarettes (vaping) were all grouped under the smoker label.

### Bulk RNA sequencing

Gene expression profiling from whole blood was performed on 317 individuals (Figure S1 and Table S2), totaling 30,795 unique transcripts. The counts were normalized using DESeq2 with default options and then mapped to the homo sapiens gene *Ensembl* database. Transcripts that could not be mapped to the database were discarded, reducing their number to 23,617. Transcripts were aggregated into Blood Transcriptome Modules (BTMs) following BloodGen3Module^44^, a fixed repertoire of 382 transcriptional modules, for patient similarity calculations. A one-dimensional Principal Component Analysis (PCA) was applied to each group of genes constituting a module and the resulting score was assigned to the corresponding BTM.

BTMs can thus be seen as metagenes, capturing variation of subsets of genes that tend to be co-expressed. Only genes included in the BTMs were considered for this aggregation.

### Blood bulk CD4+ and CD8+ TCR sequencing

The individuals for which bulk TCR sequencing data was available were 99 (Figure S1 and Table S2). Across the cohort, the number of unique CDR3 sequences were 2,024,226. To significantly reduce the feature space, and to follow the strategy chosen for RNAseq, the sequences were aggregated in 999 pre-identified co-expression TCR modules. These modules were constructed using the ImmuneCODE database^45^. Our approach focused on public clonotypes, defined as unique combinations of V-gene and TCR CDR3 amino acid sequences shared across 20 or more of the 1,468 repertoires in the dataset. For each of these clonotypes, we created a 1,486-dimensional binary vector indicating its presence or absence in each repertoire. To measure co-occurrence similarity of two clonotypes, we calculated the Pearson correlation between their binary vectors. We then constructed a k-nearest neighbor graph, connecting each clonotype with its 200 most similar neighbors. Finally, we extracted the 999 co-expression modules using spectral modularity maximization clustering of this network, implemented through the cugraph library. The value assigned to each module corresponded to the fraction of sequences relative to the patient’s repertoire size that belonged to said module.

Six statistics were used for testing to compare clusters in the downstream analysis. *TCR Covid match CD4* and *TCR Covid match CD8* represent the ratio of overlapping unique sequences of the CD4+ and CD8+ repertoires with curated SARS-CoV-2-specific TCRs of the ImmuneCODE database^45^, respectively. *TCR Covid freq CD4* and *TCR Covid freq CD8* represent the sum of reads for those TCRs identified as SARS-CoV-2-reactive, normalized by the repertoire size. Lastly, *entropy CD4* and *entropy CD8*, denote the normalized Shannon entropy for the CD4+ and CD8+ compartments.

### Microbiome

Stool samples were collected for 182 individuals (Figure S1 and Table S2). Sequencing of the highly conserved 16s RNA region was performed to characterize the bacterial strains. To address taxonomic heterogeneity, causing zero-inflation^46^, only bacteria present in at least 10% of the individuals (maximum 90% of missing values) were considered as input to calculate patient similarity. Thus, the number of different genera selected amounted to 387. Consistently with the previous two modalities, a dimensionality reduction was applied using a predefined set of eigentaxa, 84 sets of bacteria that were seen to be co-expressed. Only 75 out of 84 had at least one strain from the preselected bacteria. Bacterial abundances were normalized with DESeq2. PCA was applied for each eigentaxa on the strains belonging to said group of bacteria. The first PC was used as the value for the eigentaxa.

### CyTOF

Blood samples of 384 patients (Figure S1 and Table S2) were collected for flow cytometry immune cell characterization through sorting of cell surface markers-specific fluorescently dyed antibodies. Percentages of 144 subpopulations with similar markers were identified among the cells and used as features to compute similarities between subjects. Statistical testing to evaluate the differences in the immune cell composition between individuals was conducted on 22 manually annotated meta features, essentially a higher-level organization of the subpopulations.

During the initial development of the analysis, 36 samples were identified to have undergone suspected degradation during cryopreservation. After a selection based on percentage of unstained cells, 109 samples were removed. The threshold of the percentage of unstained cells was chosen to be of maximum 10%. After preprocessing, the remaining 236 were used to investigate similarities.

### Data integration

Patient similarity matrices were built using snfpy (v 0.2.2.)^47^. For the CyTOF and RNAseq datasets, the default squared Euclidean distance was the metric of choice, while for the microbiome and TCR modalities, the Bray-Curtis distance was used^48^ due to its biological rationale. All data was normalized prior to integration to reduce the effect of different scales. During multimodal analysis, affinity matrices from different modalities were combined using Similarity Network Fusion (SNF)^18^. SNF iteratively updates patient similarity per data type with the average similarity across all other modalities, until convergence. As proven by the original authors, SNF is robust to noise and data heterogeneity but cannot handle missing data in the chosen implementation^47^. Hence, to maximize the sample size and the power of our analysis, the unimodal affinity matrices, built using all available patients, were the starting point for multimodal integrations. Samples lacking at least one modality were only removed prior to SNF.

### Clustering

The cluster individuals, spectral clustering^49^ was chosen due to its specificity for similarity matrices. The implementation used for spectral clustering (scikit-learn v 1.2.1) initially constructs the normalized Laplacian of the affinity matrix. The first *k* largest eigenvectors of the Laplacian are then computed and used as features for a new matrix. K-means clustering is then applied for the final cluster assignment. This method is particularly useful when clusters are non-convex in the original space but can be separated in the new space defined by the eigen-decomposition of the graph.

Following the eigengap heuristic, the largest difference between eigenvalues sorted in descending order suggests the number of clusters. This approach is normally used when dealing with factor analysis or PCA analysis since the eigenvectors corresponding to the largest eigenvalues inform the direction of most variability in the data^50^. However, in this case, we looked at the smallest eigenvalues, particularly the second smallest, to assess graph connectedness^51^. In the ideal case scenario of *n* disconnected subgraphs, the eigenvalue zero is expected to show multiplicity *n*. However, we did not expect completely distinct clusters, as at least partial overlapping groups are to be expected in the data at hand^52^. The smaller the eigenvalues after zero, the more disconnected the existing subgraphs described by the Laplacian are.

We calculated silhouette scores to assess clusters coherence^53^ when varying parameters, namely: the size of the neighborhood used to construct the similarity matrix (from 5% to 15% of the total size of the population), the number of clusters (between 2 and 8) and the distance metric (among Euclidean, Squared Euclidean and Bray-Curtis distance). This analysis aided in the choice of best distance metrics to use for each data type. The remaining two parameters (size of neighborhood and number of clusters) were less impactful in the decrease of silhouette, albeit still inversely proportional to the scores.

### Stability analysis

Stability analysis was performed on the RNAseq modality. The goal was to study the robustness of this modality by quantifying severe outliers that, over hundreds of iterations, did not co-cluster with other samples (Figure S6A and S6B).

The samples underwent ten-fold clustering partitioning repeated 1000 times with various initializations. During each iteration, a clustering was constructed using nine folds, ensuring each sample appeared in 90% of the total runs. The co-clustering probability, representing the frequency of two samples being co-clustered across runs, was calculated (Figure S6C). Spectral clustering was then applied to these probabilities, with an additional cluster introduced to identify outliers. However, due to the nature of spectral clustering, the algorithm will always identify the chosen number of clusters, regardless of the true separation between samples (Figure S6D and S6E).

To address this analysis from another perspective, we determined the stability of the samples’ co-clustering based on the distance from any centroid. MultiDimensionalScaling was applied to the co-clustering probabilities to reduce the matrix to a lower dimensional space. A KMeans algorithm was then trained on the two-dimensional space and samples that were at a threshold distance from any centroid were considered unstable. Compared to the original estimation of unstable samples, we can see that these are more spread out across clusters (Figure S6D and S6F).

### View-specific analysis (One vs Rest)

Every analysis had a “one vs rest” setup, thus comparing all members of a cluster to the remaining individuals. An overview of significant results was finally visualized, here referred as “spider plots”.

### Logistic Regressions

Logistic regression, using as input the metadata and as predicted output the binary cluster membership (1 when the individual belonged to the cluster and 0 otherwise). Numerical data was normalized, and categorical data was one hot encoded, with one category being excluded and used as reference to avoid multicollinearity. P-values of the regression coefficients were Bonferroni corrected for multiple testing. Due to the high number of missing values (see Table S2), BAI, BDI scores and Covid infection data, even if collected, were excluded from the regression analysis. The following model was fit:

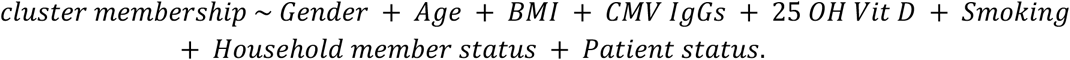

Where relevant, results of logistic regression were shows in the respective figures (Figure 1, 2, 3, 5, 6 and 7, and Figure S3 and S4).

### Mann-Whitney (MW) statistical testing

For the TCR-seq and CyTOF metafeatures (6 and 22 respectively), two-sided pairwise Mann-Whitney U statistical tests were performed. Every metafeature was compared and tested against the remainder of the cohort.

### GSEA (Gene Set Enrichment Analysis)

The number of differentially expressed genes between a cluster and the background would have made the interpretation of gene expression difficult. Compared to a single-gene analysis, Gene Set Enrichment Analysis (GSEA) softer thresholding allows for a better assessment of small signals deriving from an inherent noisy data type. GSEA was employed to mainly identify enriched gene sets from the Molecular Signatures Database (MSigDB) hallmark gene set collection^54^ but also from the Gene Ontology Biological Process, Cellular Component and Molecular Function collections. DESeq2 normalized genes were fed to GSEA to inspect if significant differences in the expression of some predetermined sets of genes sharing a common factor could be observed. Co-expression of genes belonging to certain pathways was deemed more interesting compared to increases (or decreases) of expression in single genes. Normalized Enrichment Scores (NES) based on the correlations with the condition (cluster) allowed us to select the top enriched (positively or negatively) sets. A standard value suggested by the original authors of 0.25 for the False Discovery Rate (FDR) was set to discard spurious enrichments. GSEA was implemented using the Python package GSEApy (v 1.0.4)^55^.

### Differential abundance analysis

Differential Abundance Analysis (DAA) was performed on the microbial data of the patients. In a standard differential expression analysis, DESeq2’s median of ratios normalization, is commonly used to address between-samples differences in sequencing depth and composition. The same normalization was also here implemented.

Yang and Chen^56^, comprehensively evaluated DAA methods on 106 real datasets. This study concluded that different methods could produce highly discordant results. The authors affirm that no single DAA tool should be applied in a real-world scenario blindly because of the unknown a-priori characteristics of the compositional data. Therefore, DAA should be performed with a variety of methods to determine a consensus and to avoid cherry-picking of favorable results.

Normalised taxa abundances were fed to 5 different methods from the ‘multidiffabundance’ R toolkit^57^ to test statistical differences in bacterial abundance between a cluster and the background. Namely: Corncob, Limma, clr, Maaslin2 and DESeq2. If at least two out of the five tools determined DA significance, thus the corresponding q-value scored below 0.05, the bacterial strain was selected. Log2FoldChanges of the abundances were also computed. The package *rpy2* (v 3.5.10), was used to interface the results yielded by the differential abundance testing in R with the remainder of the testing results generated in Python.

For visualization purposes, the single ASVs were here not considered, due to high clutter and lack of interpretability (16s RNAseq). However, ASVs with adjusted p-values smaller than 0.05 and absolute Log2FoldChange greater than one were still reported alongside the rest of the results and code (see Figure S2C).

### Bacterial family enrichment analysis

Investigation of existing global patterns and shared enterotypes was performed through the GSEA statistical testing framework. The same philosophy utilized for RNAseq was closely followed to test microbial differential expression at the hierarchical family level. The NES produced by GSEA was used to determine directionality and size of the family-specific effect. For this purpose, bacterial families were considered equivalent to gene sets (see Figure S2B).

Alpha diversity (Shannon entropy, Simpson Index, Chao1 richness) of the bacterial abundances were also calculated and compared between clusters. To determine statistical significance, Kruskal-Wallis tests, followed by Bonferroni-corrected Dunn’s pairwise post-hoc tests, were set in place.

### Spider plot visualizations

Sparsification of the significant results was applied to every level of the analysis. This was done to reduce visual clutter by plotting only the starkest significant differences in our spider plots. For some of the analysis, multiple testing correction allowed to filter the number of significant hits that had to be plotted, while for some other levels a threshold was established on the Log2FoldChange, filtering out results with smaller effect sizes.

It can be argued that with the (over)correction for multiple hypotheses testing we may be hindering the amount of signal we can observe. The statistical power of the testing may be reduced with this very conservative procedure. Alternative solutions such as the harmonic mean p-values for dependent tests^58^ have been proposed.

Only genesets that presented an FDR smaller than 0.25, were considered for visualization purposes. At most the first ten positively and ten negatively enriched genesets, ordered by NES, were plotted. The graph edges were weighted to represent NES, regression coefficients or Log2FCs and the color of the edge indicated the sign of enrichment. One issue that should be highlighted is that these three measures are not fully comparable since they are very different statistics. The LR coefficients represent the change in log odds generated by a unit change in the variable. The NES is derived by the sum of correlations between genes in a gene set and the phenotype, normalized by the size of the gene set or bacterial family. Finally, the Log2FoldChange is the ratio of the mean expression of the variable in the cluster of interest over the mean expression of the variable in the background, in a log2 scale.

## Acknowledgements & Funding

We appreciate the participation of all participants, patients and their families in this study. We are grateful to all unmentioned clinicians, nurses, and lab colleagues. We thank Prof. Susan Schlenner, Pier Andrée Penttila, Reena Chinnaraj, and other staff at KU Leuven Flow and Mass Cytometry Facility for their help with data acquisition on the Helios instrument. We thank dr Sofie Van Gassen and Prof Yvan Saeys for their help with FlowSOM. We would also like to thank Joke Vereecken, Maria Matteijssens, and Sam Van Goethem for their help in participant recruitment. Data analysis was performed using the resources and services provided by the Flemish Supercomputer Center, funded by the Research Foundation Flanders and the Flemish Government.

The following founding sources are acknowledged: Research Foundation Flanders (FWO) 1SH3924N (FA), 1861219N (BO), G0G4220N (KKA), G0H4520N (BO, PB, SC, RN, PVD, LK, PM, EL, EV, HG, KA, KV). European Union’s Horizon 2020 research and innovation programme grant agreement 851752-CELLULO-EPI (B.O.).

## Author contributions

Conceptualization: BO

Design: FA, MKH, TG, IDB, EB, SL, PM, BO

Experiments: FA, MKH, IDB, MK, RV, VDV, HJ, HDR, JS, KP, AS

Visualization: FA

Funding acquisition: SC, RN, KKA, KV, EV, PB, PVD, HG, EL, KL, PM, BO

Project administration: BO

Supervision: EB, SL, PM, BO

Writing – original draft: FA, PM, BO

Writing – review & editing: all authors

## Data availability

All data and code utilized for the study are available at https://github.com/fabio-affaticati/activ_covid-tcell-omics.

