## Supplementary Material for "Bridging immunotypes and enterotypes using a systems immunology approach"

### Supplementary Materials

Figures S1 to S6
Tables S1 to S2.

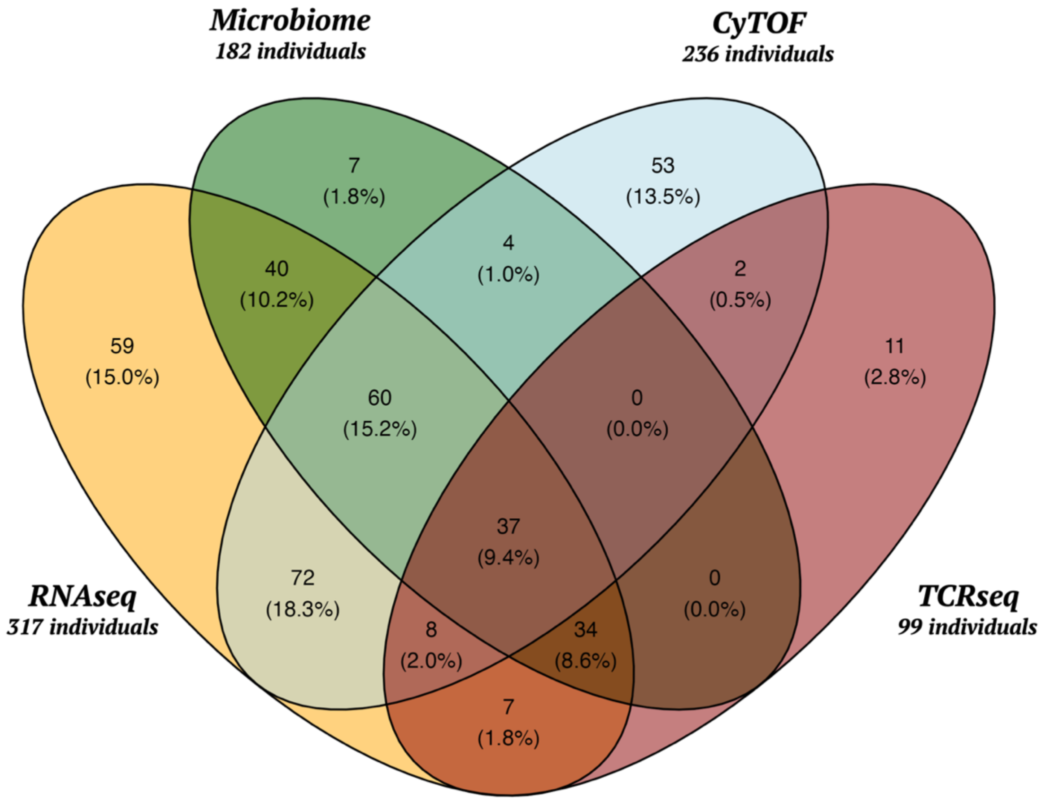


*Figure S1: Visual partitioning of the study cohort between its datatypes.*
*Venn diagram representing the distribution of the data, subdivided in the available modalities. In each section the number indicating the samples available for the intersection of modalities is shown and, between parentheses, the percentage of the total cohort thus included. The number of patients for each datatype are also shown in bold.*

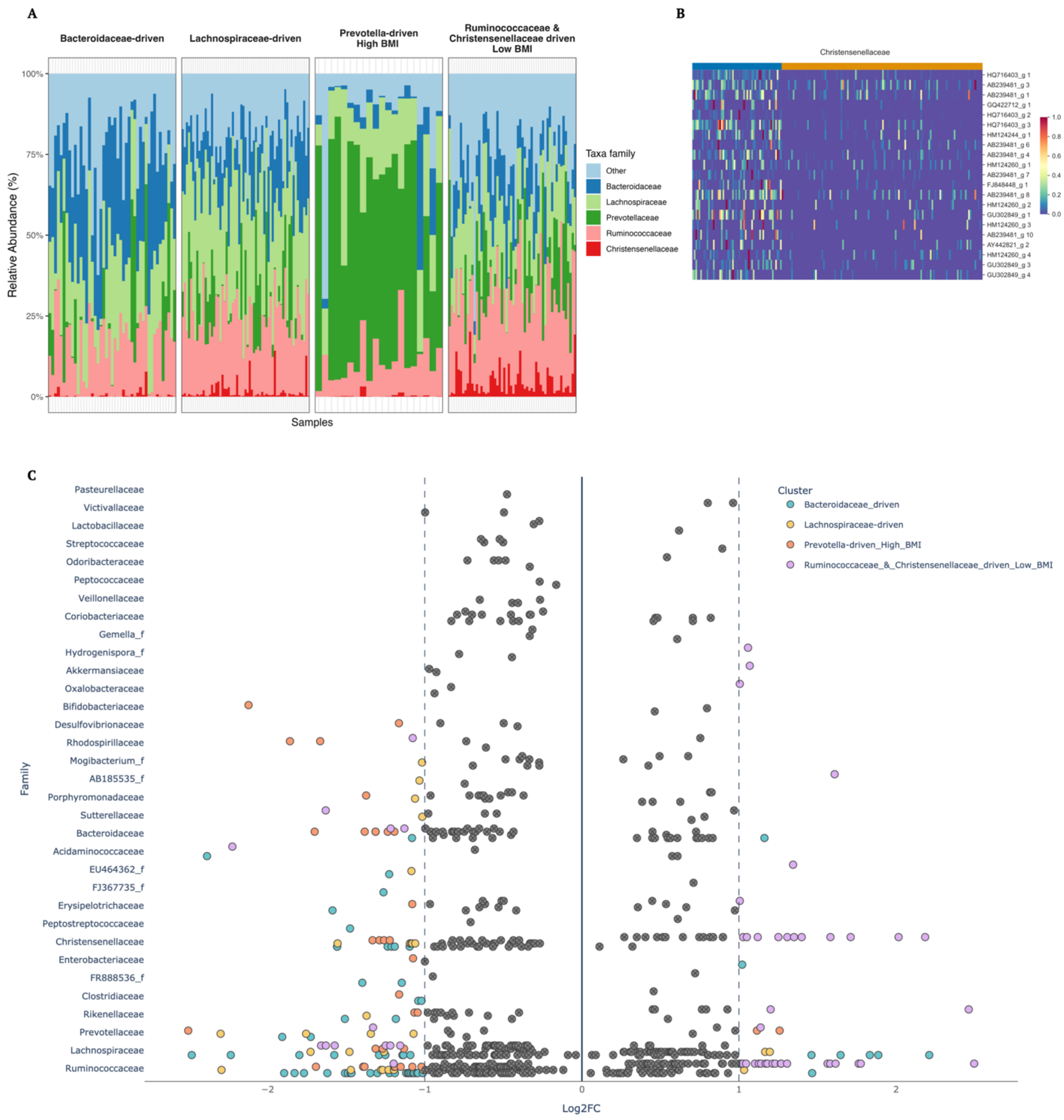


*Figure S2: Microbiome unimodal taxonomic profile.*

*(A) Relative abundances bar plot highlights sharp differences between the four microbiome unimodal clusters.*

*(B) Example of a comprehensive abundance enrichment at the family level. Christensenellaceae are here clearly shown to be overabundant in the Ruminococcaceae & Christensenellaceae driven cluster, represented by the blue annotation at the top of the plot, against, in yellow, the remainder of the cohort.*

*(C) Example of bacterial differential abundance analysis results performed on the gut microbiome normalized abundances of the cohort. ASVs with adjusted p-values smaller than 0.05 are reported. Furthermore, ASVs with absolute Log2FoldChange smaller than one are grayed out. For visualization purposes and lack of interpretability, the single ASVs were not considered in the main analysis.*

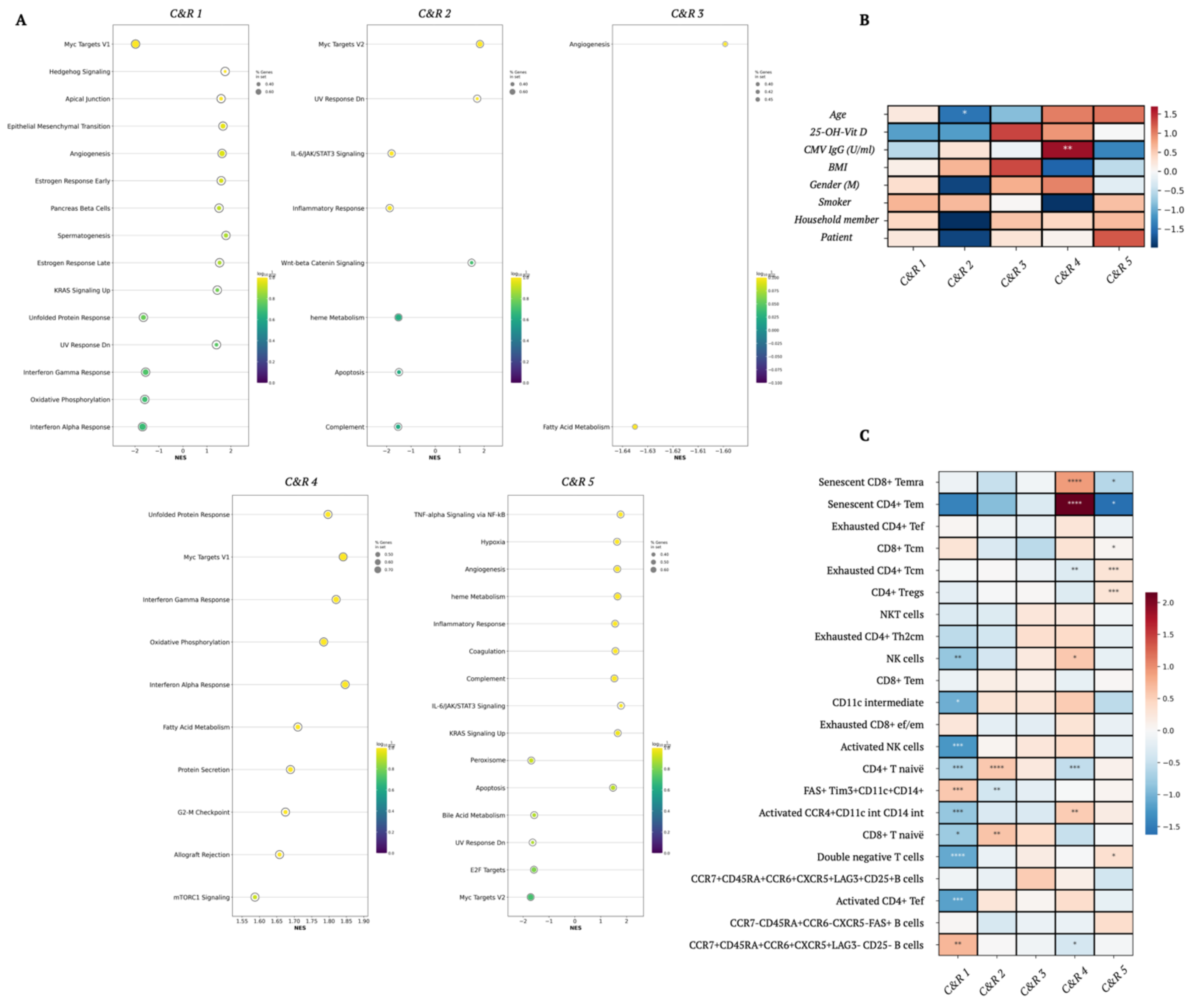


*Figure S3: The blood transcriptome captures orthogonal patterns from the gut microbiome and immune cell composition.*

*(A) Gene Set Enrichment Analysis (GSEA) Normalized Enrichment Scores (NES) in a one-vs-rest setup for the bimodal CyTOF & RNAseq case (n=177). Only gene sets enriched at a False Discovery Rate (FDR) < 0.25 are here reported. The hue represents the FDR while the dot size shows the ratio of genes in the gene set after filtering out those genes not in the expression dataset.*

*(B) Results of a logistic regression (LR) on the metadatafor the bimodal CyTOF & RNAseq case. The hue indicates the LR coefficients obtained by comparing each cluster against the background cohort. Statistical significance of the coefficients is indicated with an asterisk notation.*

*(C) Heatmap of the statistical testing results contrasting 22 cytometry patterns (metafeatures) for each patient cluster against the remainder of the cohort. The hue indicates, for each feature, the log2 ratio of its mean expression in the cluster over the mean expression in the background. P-values were obtained from Mann-Whitney u tests assessing significance adjusted for multiple testing with the Bonferroni method.*

*Asterisks indicate statistical significance ($*p < 0.05$, $**p < 0.01$, $***p < 0.001$, and $****\ p < 0.0001$) and non-significance when missing.*

*Abbreviations: Temra, terminal effector memory T cells; Tem, effector memory T cells; Tef, effector memory T cells; Tcm, central memory T cells; NK, natural killer cells; NKT, natural killer T cells; int, intermediate; C&R, CyTOF and RNAseq.*

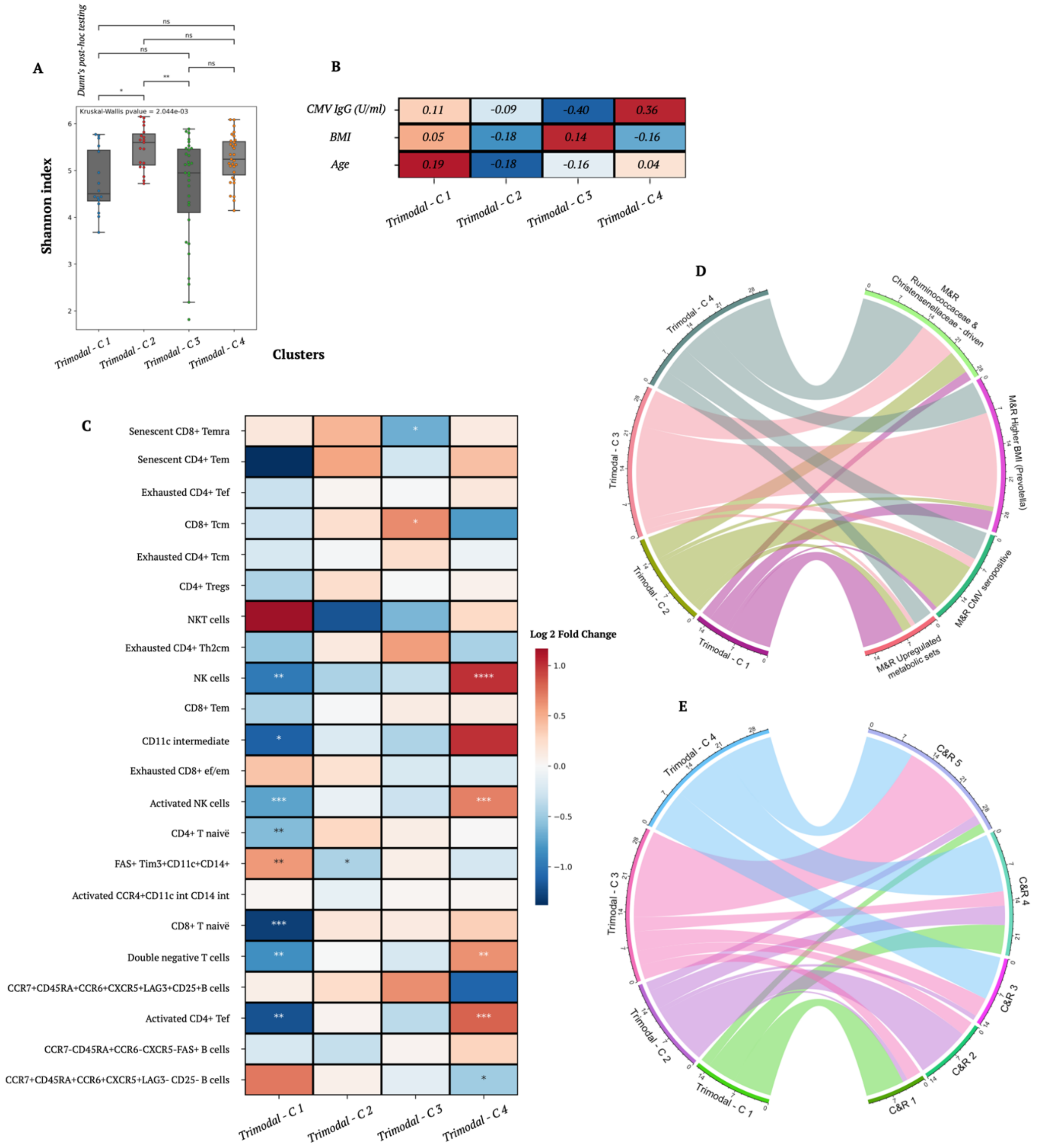


*Figure S4: Trimodal integration of RNAseq, Microbiome and CyTOF.*

*(A) Shannon index indicating intra-cluster alpha diversity of the trimodal integration of RNAseq, Microbiome and CyTOF (n=97). Kruskal-Wallis test's p-value is reported on the plot while Bonferroni corrected Dunn's post hoc tests statistical significance is indicated with asterisks.*

*(B) Results of logistic regression on the metadata for the trimodal clusters. Values of the coefficients are reported, standard significance (0.05) after correction was not reached by any feature.*

*(C) Heatmap of the statistical testing results contrasting 22 cytometry patterns (metafeatures) for each patient cluster against the remainder of the cohort. The hue indicates, for each feature, the log2 ratio of its mean expression in the cluster over the mean expression in the background. P-values were obtained from Mann-Whitney u tests assessing significance adjusted for multiple testing with the Bonferroni method.*

*Asterisks indicate statistical significance ($*p < 0.05$, $**p < 0.01$, $***p < 0.001$, and $****p < 0.0001$) and non-significance when missing.*

*(D) Partition of the CyTOF, Microbiome and RNAseq trimodal clustering (left) in the bimodal RNAseq and Microbiome clusters (right) comparing consistency.*

*(E) Partition of the CyTOF, Microbiome and RNAseq trimodal clustering (left) in the bimodal RNAseq and CyTOF clusters (right) comparing consistency.*

*Abbreviations: Temra, terminal effector memory T cells; Tem, effector memory T cells; Tef, effector memory T cells; Tcm, central memory T cells; NK, natural killer cells; NKT, natural killer T cells; int, intermediate.*

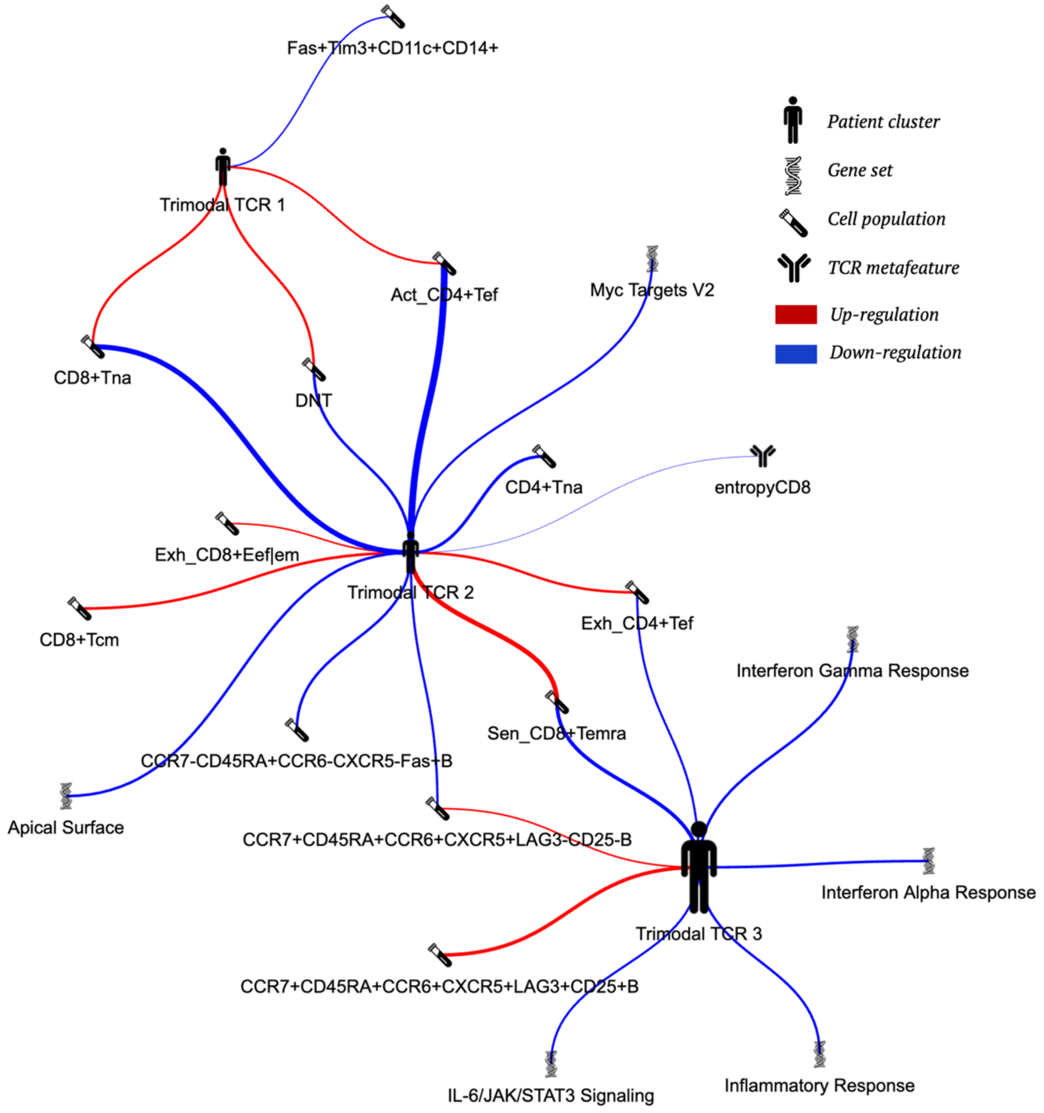


*Figure S5: Data integration across all layers is driven by inflammation markers.*

*Spider plot drawing a summary of the trimodal integration of RNAseq, CyTOF and TCRseq. Only significant results for the respective tests are reported. Red and blue arches indicate significantly positive and negative regulation of the corresponding variable in the cluster. The arch width has been scaled per analysis and indicates the effect size.*

*Abbreviations: Exh, exhausted; Sen, senescent; Act, activated; na, naïve; Tem, effector memory T cells; Tef, effector memory T cells; Tcm, central memory T cells; DNT, double negative T cells; int, intermediate.*

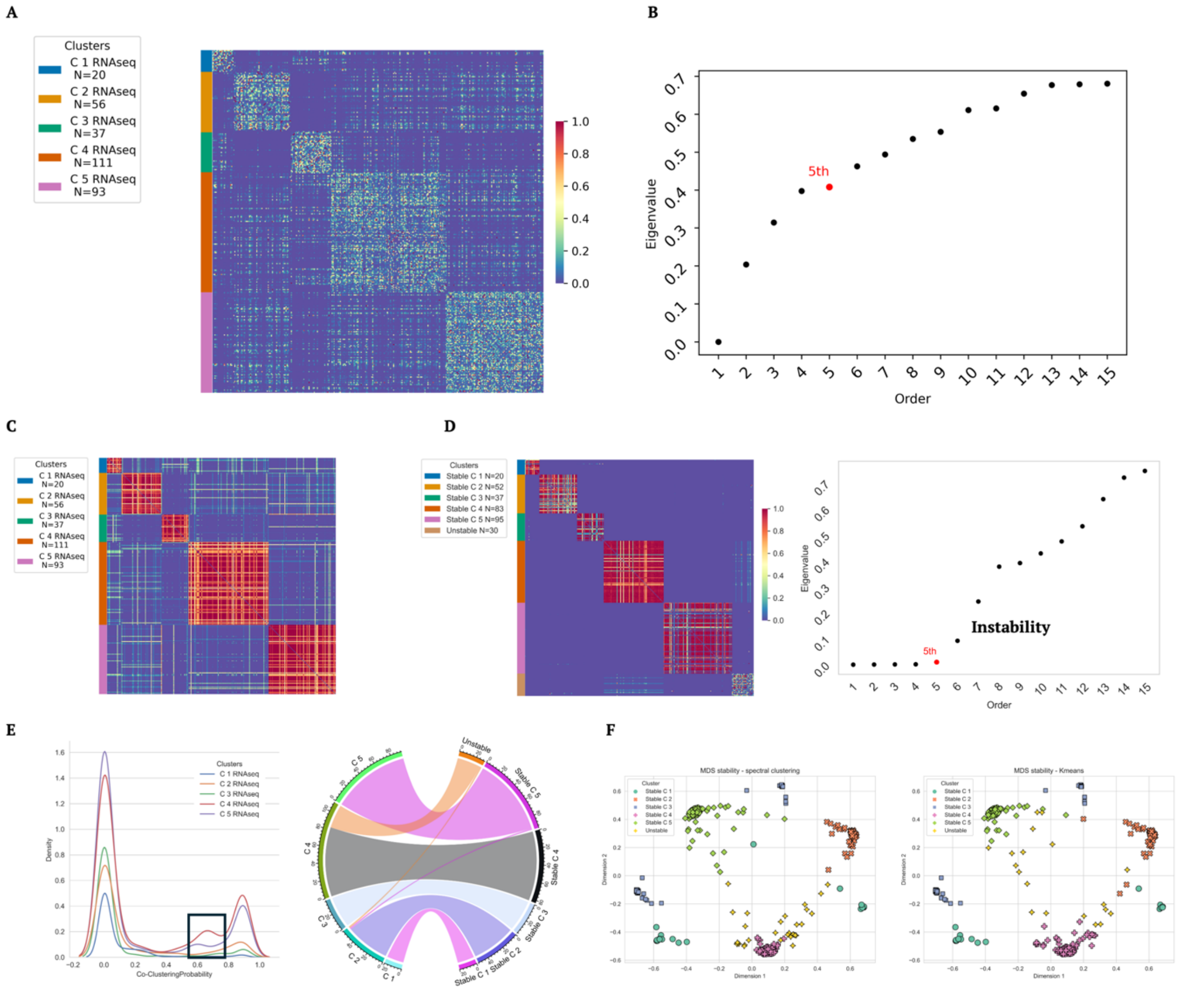


*Figure S6: Step-by-step results of the RNAseq unimodal stability analysis.*

*(A) Squared Euclidean distance matrix for the RNAseq modality. Values were minmax-scaled post-hoc to enhance the color contrast for visualization purposes without affecting the clustering process.*

*(B) The top 15 eigenvalues of the Laplacian matrix sorted in ascending order to highlight the connectivity profile of the underlying data. The fifth eigenvalue is highlighted based on the largest gap with the following eigenvalue, thus determining the number of chosen clusters. Choosing five clusters leads to a silhouette score of 0.55.*

*(C) Heatmap representing the co-clustering probabilities over the 1000 runs of. During each iteration, a clustering was constructed using nine folds, ensuring each sample appeared in 90% of the total runs. The co-clustering probability, representing the frequency of two samples being co-clustered across runs, was calculated.*

*(D) Spectral clustering was then applied to these probabilities, with an additional, $6^{th}$, cluster introduced to identify outliers. The unstable cluster silhouette score resulted in a low 0.12.*

*(E) The largest number of "unstable" samples come from a specific cluster (Cluster 4). However, due to the nature of spectral clustering, this algorithm will always identify the chosen number of clusters, regardless of the true separation between samples.*

*(F) Original estimation of unstable samples (left) compared to the Kmeans clustering in the MultiDimensionalScaling two-dimensional space (right).*

| ***CyTOF***<br>(236) | ***Microbiome***<br>***(182)*** | ***TCRseq***<br>***(99)*** | ***Microbiome & RNAseq***<br>**(171)** | ***CyTOF & RNAseq***<br>***(177)*** | ***RNAseq & Microbiome & CyTOF***<br>**(97)** | ***RNAseq & TCRseq***<br>**(86)** | ***TCRseq & CyTOF***<br>**(47)** | ***RNAseq & CyTOF & TCRseq***<br>**(45)** |
|---|---|---|---|---|---|---|---|---|
| LAG3 cluster<br>(4) [1.7%] | Prevotella driven-High BMI<br>(20) [11%] | TCR C1<br>(16) [16.2%] | M&R upregulated metabolic sets<br>(34) [19.9%] | C&R 1<br>(20) [11.3%] | Trimodal C1<br>(16) [16.5%] | Non-patient T&R cluster<br>(20) [23.3%] | T&C C1<br>(13) [27.7%] | Trimodal TCR 1<br>(10) [22.2%] |
| Monocyte-driven<br>(11) [4.7%] | Bacteroidaceae-driven<br>(45) [24.7%] | TCR C2<br>(6) [6.1%] | M&R CMV seropositive<br>(36) [21.1%] | C&R 2<br>(28) [15.8%] | Trimodal C2<br>(19) [19.5%] | Non-Covid T&R inflammation cluster<br>(16) [18.6%] | T&C C2<br>(16) [34%] | Trimodal TCR 2<br>(11) [24.4%] |
| CMV seropositive senescent<br>(56) [23.7%] | Ruminococcaceae & Christensenellaceae driven-Low BMI<br>(56) [30.8%] | TCR C3<br>(41) [41.4%] | M&R higher BMI (Prevotella)<br>(45) [26.3%] | C&R 3<br>(29) [16.4%] | Trimodal C3<br>(31) [32%] | T&R cluster 3<br>(20) [23.3%] | T&C C3<br>(18) [38.3%] | Trimodal TCR 3<br>(24) [53.3%] |
| FAS cluster<br>(44) [18.6%] | Lachnospiraceae-driven<br>(61) [33.5%] | TCR C4<br>(36) [36.4%] | M&R Ruminococcaceae & Christensenellaceae driven<br>(56) [32.7%] | C&R 4<br>(44) [24.9%] | Trimodal C4<br>(31) [32%] | Covid T&R inflammation cluster<br>(30) [34.9%] | | |
| Naive group<br>(57) [24.2%] | | | | C&R 5<br>(56) [31.6%] | | | | |
| Elderly group<br>(64) [27.1%] | | | | | | | | |

*Table S1: Summary of the cluster sizes across all analyses.*

*Columns represent each analysis, named after the feature spaces included, along with the total number of individuals included. In each column, representative names for each patient cluster identified are listed. Between parenthesis the size in number of individuals and between squared brackets the size percentage.*

*Abbreviations: CMV, cytomegalovirus; BMI, Body-Mass Index; M&R, Microbiome and RNAseq; C&R, CyTOF and RNAseq; T&R, TCRseq and RNAseq; T&C, TCRseq and CyTOF.*

| *Data type* | *CyTOF* | *Microbiome* | *RNAseq* | *TCRseq* | *Metadata* |
|---|---|---|---|---|---|
| *Subject count* | *236* | *182* | *317* | *99* | *569* |
| *Gender (male/female)* | 91/152 (1%) | 81/96 (3%) | 136/167 (4%) | 44/45 (10%) | 213/321 (6%) |
| *Age (years)* | 51 [33.5-61] (2%) | 56 [46-64] (3%) | 54 [38.25-63] (5%) | 60 [50-66] (10%) | 49.5 [34.75-60] (1%) |
| *BMI (kg/$m^2$)* | 25.51 [23.03-28.72] (1%) | 26.09 [23.27-29.32] (3%) | 25.99 [23.3-29.1] (4%) | 27.04 [24.62-30.32] (10%) | 25.25 [22.68-28.62] (6%) |
| *CMV IgG (U/ml)* | 0.15 [0.15-262.5] (7%) | 0.15 [0.15-166] (8%) | 0.15 [0.15-228] (9%) | 0.15 [0.15-284] (14%) | 0.15 [0.15-271.5] (10%) |
| *25-OH Vitamin D (ng/ml)* | 20 [14-27] (7%) | 20 [16-28] (8%) | 20 [16-28] (9%) | 21 [16-30] (14%) | 20 [15-27] (10%) |
| *Smoker (yes/no)* | 23/210 (1%) | 9/168 (3%) | 24/279 (4%) | 4/85 (10%) | 55/478 (6%) |
| *Group membership (control/patient/household member)* | 154/65/14 (1%) | 102/66/9 (3%) | 165/117/21 (4%) | 58/16/15 (10%) | 254/164/151 (0%) |
| *BDI T-score depression* | 54 [47.25-60.75] (74%) | 53 [44-58] (71%) | 53.5 [44-59] (74%) | 47 [42-58] (78%) | 53 [44-58] (76%) |
| *BAI T-score anxiety* | 51 [47-59.75] (74%) | 51 [46-55] (71%) | 51 [46-57] (74%) | 51.5 [44-56.5] (78%) | 51 [46-59] (74%) |
| *Covid infected (yes/no)* | 65/165 (3%) | 67/108 (4%) | 117/182 (6%) | 58/28 (13%) | 208/297 (11%) |
| *Long Covid symptoms (yes/no)* | 35/30 (0%) | 31/36 (0%) | 57/60 (0%) | 31/27 (0%) | 91/117 (0%) |
| *TCR Covid freq CD8* | - | - | - | 4.9 * 10^-4 [0.6 * 10^-4 – 8.3 * 10^-4] | - |
| *TCR Covid freq CD4* | - | - | - | 5.6 * 10^-4 [ 1.1 * 10^-4 – 8.0 * 10^-4] | - |
| *TCR Covid match CD8* | - | - | - | 1.1* 10^-3 [2.9 * 10^-4 – 1.4 * 10^-3] | - |
| *TCR Covid match CD4* | - | - | - | 6.6 * 10^-4 [3.0 * 10 ^-4 - 8.8 * 10^-4] | - |
| *Entropy CD8* | - | - | - | 0.81 [0.75-0.85] | - |
| *Entropy CD4* | - | - | - | 0.93 [0.83-0.95] | - |

| | | | | | |
|---|---|---|---|---|---|
| % Senescent CD8+ Temra | 4.78 [2.2-8.54] | - | - | - | - |
| % Senescent CD4+ Tem | 0.28 [0.06-2.02] | - | - | - | - |
| % Exhausted CD4+ Tef | 2.08 [1.59-2.82] | - | - | - | - |
| % CD8+ Tcm | 0.49 [0.26-0.85] | - | - | - | - |
| % Exhausted CD4+ Tcm | 15.4 [12.17-18.77] | - | - | - | - |
| % CD4+ Tregs | 0.99 [0.75-1.26] | - | - | - | - |
| % NKT | 0.81 [0.43-1.66] | - | - | - | - |
| % Exhausted CD4+ Th2cm | 0.51 [0.34-0.75] | - | - | - | - |
| % NK | 7.05 [3.8-9.83] | - | - | - | - |
| % CD8+ Tem | 2.62 [1.37-4.09] | - | - | - | - |
| % CD11c int | 0.29 [0.18-0.47] | - | - | - | - |
| % Exhausted CD8+Eef/em | 5.04 [3.56-7.33] | - | - | - | - |
| % Activated NK | 0.16 [0.07-0.31] | - | - | - | - |
| % Naïve CD4+ | 16.92 [11.86-21.62] | - | - | - | - |
| % Naïve CD8+ | 4.91 [2.23-7.48] | - | - | - | - |
| % Fas3+Tim3+CD11 | 15.35 [12.06-20.04] | - | - | - | - |
| % Activated CCR4+CD11c int CD14 int | 0.087 [0.049-0.138] | - | - | - | - |
| % DNT | 0.055 [0.032-0.089] | - | - | - | - |
| % CCR7+CD45RA+CCR6+CXCR5+LAG3+CD25+ B cells | 0.131 [0.091-0.205] | - | - | - | - |
| % Activated CD4+ Tef | 0.160 [0.055-0.270] | - | - | - | - |
| % CCR7-CD45RA+CCR6-CXCR4-Fas+ B cells | 0.064 [0.038-0.093] | - | - | - | - |
| % CCR7+CD45RA+CCR6+CXCR5+LAG3-CD25- B cells | 7.49 [5.52-9.51] | - | - | - | - |

*Table S2: Cohort characteristics.*

*Demographics data, clinical data and (when relevant for the data type) peripheral immune cell counts and TCR summary statistics are reported.*
*Numerical data is represented as median [25$^{th}$ percentile – 75$^{th}$ percentile], while categorical data category sizes are shown separated by '/'. Between parenthesis the percentage of missing values.*

*Abbreviations: CMV, cytomegalovirus; BMI, Body-Mass Index; BDI T-score , Beck Depression Inventory score; BAI T-score , Beck Anxiety Inventory score; Temra, terminal effector memory T cells; Tem, effector memory T cells; Tef, effector memory T cells; Tcm, central memory T cells; NK, natural killer cells; NKT, natural killer T cells; int, intermediate; DNT, Double Negative T cells.*
